# A translational checkpoint couples proline sensing to mitochondrial proline catabolism in *Candida glabrata*

**DOI:** 10.64898/2026.03.05.709770

**Authors:** Aishwarya Rana, Shirish Kumar Gangber, Anjli Tanwar, Anil Thakur

**Affiliations:** Regional Centre for Biotechnology, 3rd Milestone Gurgaon-Faridabad Expressway Faridabad 121001, India

**Keywords:** Fungal pathogens, *Candida glabrata*, proline catabolism, Gcn2–Gcn4, *PUT3*, metabolism, translation regulation

## Abstract

Proline catabolism represents a central metabolic and regulatory hub integrating nutrient sensing, stress adaptation, and energy production across diverse organisms. Despite its biological importance, the regulatory mechanisms controlling proline catabolism remain poorly understood in eukaryotic microbes. Here, we describe the transcriptional and translational coordination to catabolize the proline to maintain the cellular homeostasis under stress in the human fungal pathogen *Candida glabrata*. We identified proline utilisation trigger global translation repression to activates the stress-sensing kinase Gcn2, which phosphorylates eIF2α, thereby promoting the activation of the transcription factor Gcn4. Activated Gcn4 upregulates the transcription factor Put3 and the proline transporter Put4. Put3 orchestrates expression of the mitochondrial catabolic enzymes Put1 and Put2, ensuring efficient proline utilization, mitochondrial function, and redox balance. Genetic disruption of *PUT3* abolishes proline utilization, impairs mitochondrial function, and severely compromises cellular fitness. Importantly, Put3-mediated proline catabolism is also critical for *C. glabrata* survival within macrophages and for virulence in systemic infection models. These findings reveal a mechanistic link between proline catabolism, translational regulation, and amino acid sensing in *C. glabrata*. We propose a regulatory cascade wherein Gcn2–Gcn4–Put3 signaling aligns translational reprogramming with metabolic demands to optimize proline utilization. Thus, this study establishes proline catabolism as a signaling-driven adaptive mechanism essential for fungal metabolism and persistence, rather than merely a nutritional pathway.

**AUTHOR SUMMARY:** Proline is a versatile amino acid that is essential for cellular metabolism, signaling, stress adaptation, and redox equilibrium. Proline catabolism has been implicated in cancer biology and is increasingly recognized as a key determinant of virulence in diverse pathogens. Although the enzymatic processes of proline use are well characterized, the regulatory mechanisms that sense proline availability and coordinate its metabolic integration remain poorly understood. Here, we identify a hitherto unknown regulatory axis linking proline catabolism to translational reprogramming in *Candida glabrata*, which seems to be similarly present in numerous fungi. We demonstrate that proline utilization triggers global translational repression via Gcn2-dependent phosphorylation of eIF2α, thereby activating the transcription factor Gcn4. Gcn4 is essential for proline utilization, as it controls the expression of *PUT3* and *PUT4*. Notably, *PUT3* has no known human counterpart. Our findings establish proline as a metabolic signal that couples translational control to virulence, revealing new opportunities for antifungal intervention.

## INTRODUCTION

Proline is a multifunctional amino acid that not only serves as a building block of proteins but also serves as an essential link between redox homeostasis, energy metabolism, stress tolerance, and lifespan (1, 2). Due to its unique cyclic structure and redox-active metabolism, proline contributes to adaptation under fluctuating nutrient and oxidative stress conditions (3). It represents a readily accessible nitrogen reservoir for pathogens within host environments, as it constitutes nearly 25% of collagen, the most abundant protein in the extracellular matrix, and is released through host proteolysis (2).

Proline metabolism serves as a signaling system regulating cellular physiology and stress adaptation in various species (4–6). Pathogens such as *Helicobacter pylori*, *Staphylococcus aureus*, and *Mycobacterium tuberculosis* exploit proline oxidation to sustain metabolism and maintain redox balance during periods of nutrient limitation or host-induced stress (5). Proline also aids fungal pathogens in carbon and nitrogen assimilation while activating stress-response pathways essential for survival in host environments (7). Thus, fungal pathogens rely heavily on metabolic flexibility, particularly on utilizing amino acids, to survive in nutrient-limited, oxidative, and immune-restrictive host conditions (8–10). Fungal pathogens can also upregulate proline catabolism in response to mucin and collagen-derived substrates, underscoring the ecological and physiological relevance of this pathway in host adaptation (7). However, while the biochemical steps of proline catabolism mediated by the mitochondrial enzymes proline oxidase (Put1) and P5C dehydrogenase (Put2) are well defined in *Saccharomyces cerevisiae* and *Candida albicans* (11, 12). The regulatory mechanism governing this pathway remains enigmatic. Understanding how eukaryotic cells sense proline availability and integrate it into broader regulatory metabolic networks remains a crucial question. Although proline catabolism has been thoroughly investigated across multiple biological systems, the upstream signals that trigger this pathway and the regulatory mechanisms that control its activation remain largely unknown.

*Candida glabrata*, an opportunistic human pathogen, exhibits enhanced drug resistance and high oxidative stress tolerance, indicative of efficient stress adaptation strategies. These features contribute to its “stealth,” “evasion,” and “persistence” behaviors, allowing it to survive and replicate within host phagosomes while minimizing immune activation and tissue damage (13–17). A crucial yet understudied determinant of *C. glabrata* persistence is metabolic flexibility, the ability to utilize a range of nitrogen and carbon sources under nutrient-limited and variable host conditions. Diverse nitrogen sources, such as amino acids (serving as both nitrogen and carbon sources), also enhance *C. glabrata* virulence by supporting biofilm formation and survival in the host (18–20). Additionally, several metabolic processes involving amino acids, particularly morphogenetic amino acids, like methionine, proline, and arginine, regulate both in vitro and in vivo fungal pathogen survival (4, 7, 21–23).

Proline is unique among amino acids due to its involvement in both metabolism and stress tolerance (1, 5, 24). Hence, the present study investigated the regulation of proline catabolism in *C. glabrata*. Despite its phylogenetic proximity to the non-pathogenic yeast *S. cerevisiae* and its inability to form true hyphae, *C. glabrata* exhibits potent virulence, as evidenced by an extensive repertoire of virulence traits, including adherence, secretion of hydrolytic and proteolytic enzymes, biofilm formation, and notable genome plasticity (25–27). We have described the regulatory mechanism which control proline utilization from sensing to catabolism. Proline utilization generates global translation repression that triggers Gcn2-dependent eIF2α phosphorylation, which selectively activates Gcn4. Activated Gcn4 then orchestrates the transcriptional activation of the proline utilization machinery by regulating the *PUT3* transcription factor and the *PUT4* proline transporter. This cascade links metabolic flux to stress adaptation by creating a regulatory loop wherein proline availability drives its own catabolic activation through translational regulation. Proline sensing is a key factor in determining metabolic fitness, as the disruption of this circuit (by *put1*Δ, *put2*Δ, or *put3*Δ) alters proline utilization and mitochondrial respiration, with *put3*Δ also contributing to host survival and virulence. Proline utilization is thus, an example of a metabolic sensing axis that combines stress adaptation, translational regulation, and nutritional availability, contributing to fungal virulence. Hence, by coupling catabolic activity to translational control, eukaryotic cells can achieve dynamic homeostasis between energy production and survival.

## RESULTS

### Proline utilization represses global translation by activating eIF2α phosphorylation

*Candida glabrata*, though phylogenetically closely related to *S. cerevisiae* than *C. albicans*, displays distinct metabolic plasticity. In particular, it can utilize proline as both nitrogen and carbon source (7). Consistent with the findings of Silao et al., *S. cerevisiae* was unable to utilize proline as the sole carbon source efficiently, and supplementation with minimal dextrose (0.1% or 0.2%) did not significantly improve growth; however, it could utilize proline as a nitrogen source (Fig. 1a & S1a-b). In contrast, *C. glabrata* efficiently utilized proline as both nitrogen and carbon source under all experimental conditions, highlighting its remarkable metabolic flexibility (Fig. 1a & S1a-b). Thus, to explore metabolic flexibility in *C. glabrata*, we examined the growth dynamics of *C. glabrata* cells on proline as the sole nitrogen source. *Candida glabrata* cells exhibited robust growth with ammonium sulfate, a preferred nitrogen source. When proline was provided as the sole nitrogen source, *C. glabrata* cells initially exhibited slower growth and a longer lag phase compared to those grown on ammonium sulfate. However, after a period of acclimatization, *C. glabrata* cells adapted and resumed adequate growth, indicating metabolic reprogramming that enabled effective proline utilization following initial adjustments (Fig. 1b). Efficient proline utilization after the acclimatization phase suggested the involvement of underlying regulatory pathways coordinating nutrient sensing and metabolic adaptation. In a previous study, we demonstrated that an initial extended lag phase in the growth curve, observed under both amino acid starvation and oxidative stress, led to global translation suppression, wherein ribosomes preferentially translated stress-response transcripts to facilitate stress acclimation (17). To examine the global translation state of *C. glabrata* with respect to proline utilization, we used polysome profiling to monitor ribosome activity and compare the concentration of mRNA bound to free ribosomes under ammonium sulfate- and proline-grown conditions at different time points. Remarkably, cells grown on proline as the nitrogen source exhibited a rapid and pronounced decrease in global translation compared to those grown on ammonium sulfate. Moreover, proline exposure resulted in a substantial accumulation of 80S monosomes, polysome runoff, and a marked reduction in the polysome-to-monosome (P/M) ratio, quantified using area under the curve [AUC ] (Fig. 1c). This global decrease in translation upon proline utilization was further validated using a puromycin incorporation assay; after 30 min of proline utilization, puromycin incorporation was nearly absent when detected using anti-puromycin antibodies. Notably, translational repression was evident as early as 30 min after proline utilization. Although the incorporation reappeared later, it remained visibly lower than the untreated control at corresponding time points (Fig. S1c). This global repression was primarily mediated through phosphorylation of eIF2α (eIF2α-P) by kinase Gcn2. To investigate this, we performed immunoblot analysis to assess the eIF2α phosphorylation level in proline-grown cells. We observed a robust increase in eIF2α phosphorylation, which increased 7.68- fold after 30 min and gradually declined to 6.68- and 5.81-fold after 120 min and 240 min, respectively (Fig 1d). However, this phosphorylation was completely abolished in the *gcn2Δ* mutant (Fig. 1e). These three time points were consistently used across experiments based on the sustained levels of eIF2α-P. Interestingly, there was a gradual attenuation of both translational repression and eIF2α-P at 240 min compared to earlier time points, suggesting that the cells were adapting to the utilization of proline as a nitrogen source. Collectively, these results demonstrate that the utilization of proline generates metabolic stress, which activates Gcn2-mediated translational regulation. Thus, proline utilization in *C. glabrata* not only supports nitrogen assimilation but also engages adaptive signaling pathways that fine-tune translation, a mechanism likely critical for survival and persistence within host niches.

**Figure 1.**
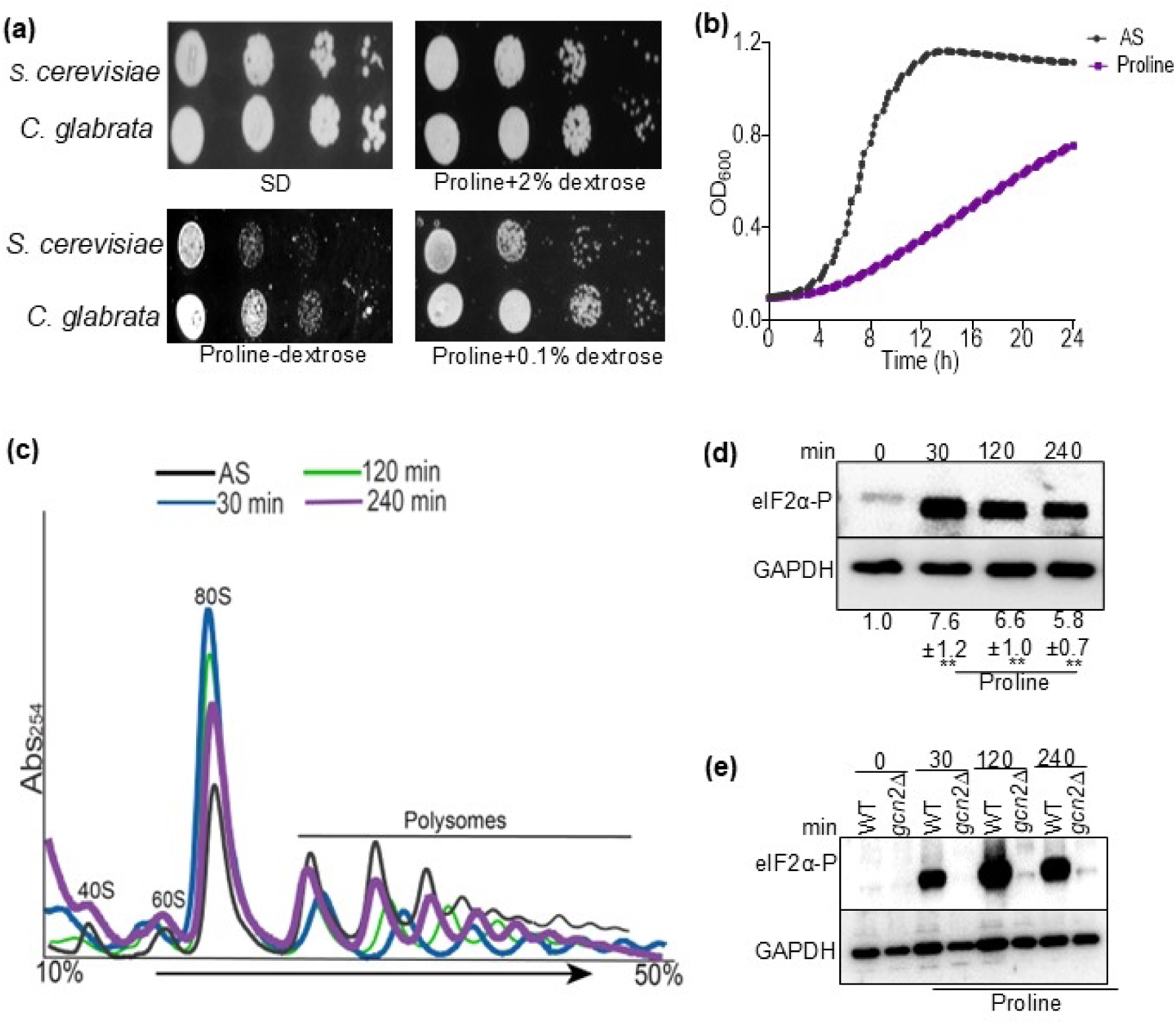
Proline utilization induces global translational repression via Gcn2-mediated phosphorylation of eIF2α. (a) Ten-fold serial dilutions of *S. cerevisiae* and *C. glabrata* cultures prepared from cells at OD_600_ ≈ 1 were spotted on control SD plates, plates supplemented with 10 mM proline, and plates containing proline with 0.1% or 2% dextrose. All plates were incubated at 30°C for 3–4 d. (b) *Candida glabrata* cells were cultivated in SD medium and in minimal media supplemented with proline as the sole nitrogen source (lacking ammonium sulfate). Growth was monitored using a Spectramax plate reader at 15-min intervals for 24 h. A representative growth curve from three independent experiments is shown. (c) *Candida glabrata* cells were grown in SD media to log phase, and then subcultured on minimal media lacking ammonium sulfate. Cells were provided proline for 30 min, 120 min, and 240 min, maintaining SD media as the control. Cells were treated with cycloheximide at various time points to arrest ribosomes. The lysates were prepared and layered onto a sucrose gradient (10–50%) to obtain profiles, and tracing was recorded at an absorbance of 254 nm. The polysome-to-monosome (P/M) ratio was quantified by calculating the AUC in ImageJ, and the mean, along with S.D., was determined from two biological replicates (n=2). d). WT *C. glabrata* cells were cultured in SD medium to log phase and transferred to minimal medium lacking ammonium sulfate to provide proline (at the indicated time points of 30 min, 120 min, and 240 min). Furthermore, WCEs (OD_600_ ∼3) were subjected to Western blot analysis using antibodies against phosphorylated eIF2α, and GAPDH was used as the loading control. eIF2α phosphorylation signals were normalized to those for GAPDH, and mean values (±S.E.M.) were calculated from three biological replicates (n=3). Statistical significance was determined using a two-tailed, unpaired Student’s *t*-test. *p<0.05, **p<0.01, ***p<0.001. (e) WT and *gcn2Δ* strains were cultured in SD medium and in minimal media containing proline as the sole nitrogen source for 30 min, 120 min, and 240 min, with an untreated SD control. WCEs (OD_600_ = ∼3) were prepared and subjected to Western blot analysis for phosphorylated eIF2α and GAPDH as described above.

### Proline utilization promotes *GCN4*-mediated translational regulation to support metabolic adaptation

Proline utilization in *C. glabrata* involves global translation repression mediated by Gcn2. Under stress conditions, Gcn2 phosphorylates the α-subunit of eIF2, a component of the ternary complex that is also associated with initiator transfer RNA (Met-tRNA_i_^Met^). This phosphorylation decreases the availability of the ternary complex, resulting in global translation inhibition. This downregulation prioritizes the selective translation of stress-responsive transcripts, including the expression of the master transcription factor *GCN4* (17, 28).

Gcn4 activates amino acid biosynthesis and general stress adaptive genes (28–32) (Fig. 2a). The Gcn2–Gcn4 pathway has been primarily associated with regulating amino acid biosynthesis, while its broader role in amino acid catabolism and metabolic adaptation remains largely unexplored. To investigate the involvement of Gcn2 in proline catabolism in *C. glabrata*, we analyzed the growth of the *gcn2*Δ strain on proline as the nitrogen source. We noted that the *gcn2Δ* mutant exhibited a pronounced growth defect, indicating the necessity of Gcn2 in proline metabolism (Fig. 2b). To determine whether Gcn2 regulates *GCN4* during proline utilization, we employed a *GCN4*-luciferase reporter construct (*GCN4*-Luc). This construct included 1000 bp of the *GCN4* 5′UTR (containing all four uORFs) cloned upstream of the *FLUC* gene (Fig. 2c) (17). When introduced into wildtype (WT) and *gcn2Δ* backgrounds, luciferase expression increased significantly at a steady pace over time (∼2.33-fold at 30 min, ∼2.87-fold at 120 min, and ∼3.71-fold at 240 min in WT upon proline utilization, whereas in *gcn2Δ* mutant, there is no significant increase at similar time points), highlighting the essential role of Gcn2 in *GCN4* activation during proline utilization (Fig. 2d). Notably, *gcn4*Δ mutant failed to grow on proline as the sole nitrogen source compared to WT and *GCN4* complemented strains, verifying the requirement of *GCN4* for proline metabolism (Fig. 2d-e). Interestingly, *GCN4*-*Luc* expression remained highly elevated in the *gcn4Δ* strain compared to WT grown on proline, indicating that the mutant strain was under stress and required *GCN4* for subsequent survival (Fig. 2f). Effectively, these results demonstrate that Gcn2-dependent activation of *GCN4* is essential for efficient proline utilization and is vital for utilizing proline as a nitrogen source.

**Figure 2.**
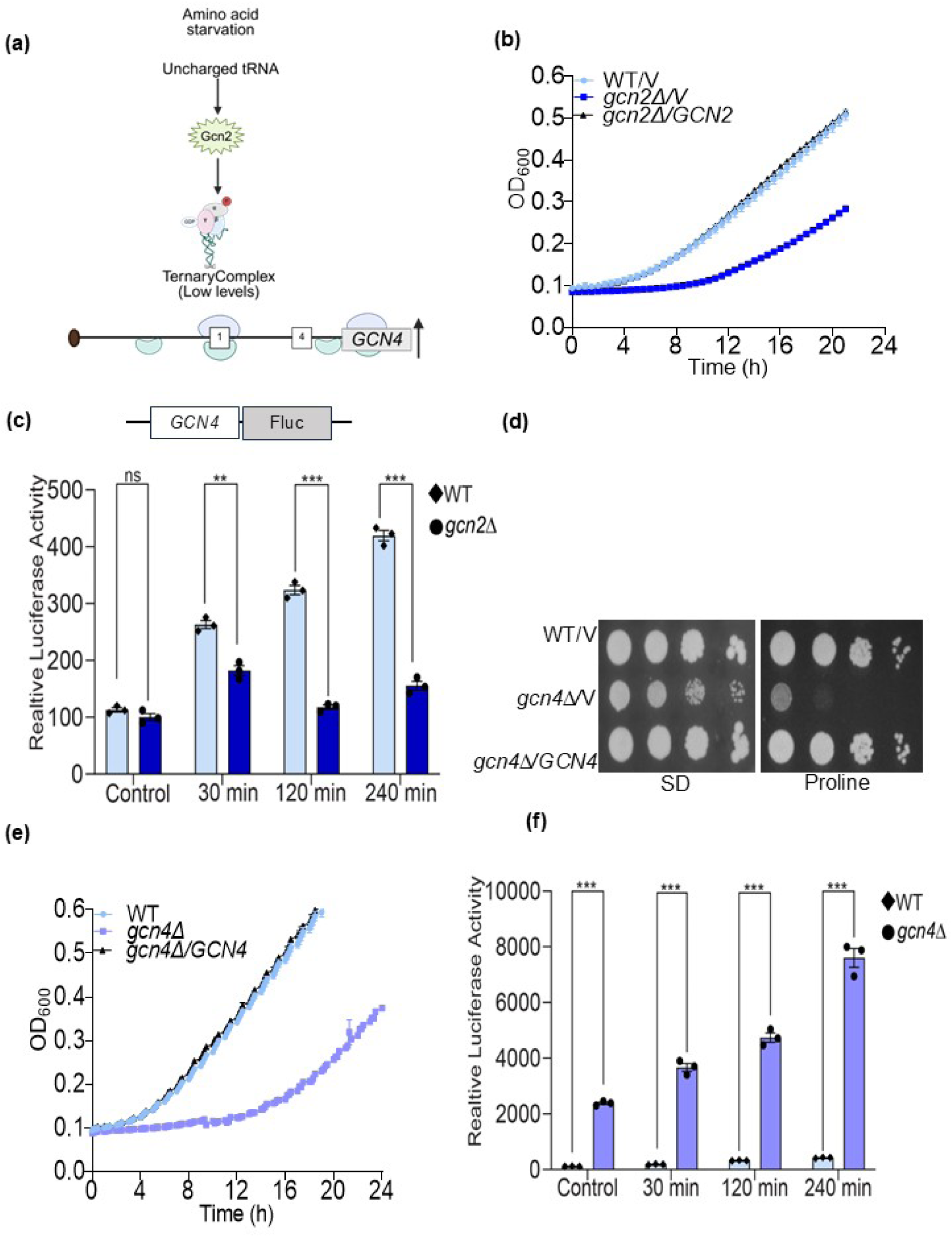
Proline utilization in *Candida glabrata* involved Gcn2-mediated *GCN4* activation. (a-b) Under unstressed conditions, the Gcn2 kinase remains inactive, the ternary complex levels are high, and the Gcn4 transcription factor remains inactive. In contrast, various stresses, including amino acid starvation, activate Gcn2, leading to eIF2α phosphorylation and a reduction in the global ternary complex pool. This results in a general downregulation of global translation but facilitates the preferential translation of the *GCN4* transcription factor, which in turn activates the expression of multiple stress-adaptive genes. (c) *Candida glabrata* WT/V, *gcn2Δ/V*, and *gcn2Δ/GCN2* strains were cultivated in SD-Ura medium and in minimal medium containing proline as the sole nitrogen source in 96-well plates. Growth was monitored using a Spectramax plate reader at 30-min intervals for 24 h. A representative growth curve from three independent experiments is shown. (d) The constructs promoter-*GCN4*-Fluc reporter and Fluc reporter (control) were transformed in WT and *gcn2Δ* strains. The cells were grown to the exponential phase and either left untreated or given proline induction for 30 min, 120 min, and 240 min, and then harvested to obtain a sample with an O.D. of approximately 1. A luminometer was used to measure luminescence, and relative luciferase units were normalized with the control reporter and total protein content (Bradford) in the samples. (e) Spot assay was performed on WT, *gcn4Δ*/V, and *gcn4Δ*/*GCN4* strains from overnight cultures in SD media. The cultures were serially diluted, spotted in SD media with ammonium sulfate and minimal media with proline as the sole nitrogen source, and incubated at 30°C for 3–4 d. (f) Growth of *C. glabrata* WT/V, *gcn4Δ/V*, and *gcn4Δ/GCN4* strains in SD-Ura medium and in minimal medium with proline as the sole nitrogen source was monitored in 96-well plates, using a Spectramax reader as described above. (f) *GCN4* expression in WT and *gcn4Δ* strains under proline induction at the previously indicated time points was measured using a luciferase reporter assay, with SD media serving as the untreated control. Promoter-*GCN4*-Fluc and Fluc reporter (control) were transformed into the Bg14 *C. glabrata* background. The relative luciferase units were normalized to the control reporter and total protein content (Bradford). The ratio of mean luciferase expression for each reporter was calculated from three biological replicates and plotted with error bars indicating S.E.M (n=3). Statistical significance was determined using a two-tailed, unpaired Student’s *t*-test. *p<0.05, **p<0.01, ***p<0.001.

### *GCN4* regulates proline catabolism by activating proline permease *PUT4* and the key transcriptional regulator *PUT3* of proline catabolic genes

Proline utilization in *C. glabrata* involves global translation repression through Gcn2-mediated eIF2α phosphorylation and subsequent activation of *GCN4*. The *gcn4*Δ mutant exhibited defective proline utilization, prompting the investigation of the underlying mechanism, with a focus on proline catabolism. This pathway comprises sequential enzymatic steps: proline oxidase (Put1) catalyzes the conversion of proline to Δ^1-pyrroline-5-carboxylate (P5C), concurrently reducing FAD and donating electrons to the mitochondrial electron transport chain, followed by Δ^1-pyrroline-5-carboxylate dehydrogenase (Put2), which converts P5C to glutamate (4, 11, 12, 24). Glutamate is then deaminated by glutamate dehydrogenase (Gdh2) to α-ketoglutarate, which enters the Krebs cycle (33, 34). Beyond this linear pathway, arginine and ornithine also contribute to the proline metabolic network, where arginine is converted to ornithine and urea via arginase (Car1), ornithine is converted to P5C by ornithine aminotransferase (Car2), and P5C can be reduced back to proline by pyrroline-5-carboxylate reductase (Pro3) (4, 35). Thus, proline serves as a central metabolic node linking multiple amino acid degradation pathways (Fig. 3a). To dissect the role of *GCN4* in proline catabolism, we evaluated the growth of WT, *gcn4Δ*, and *gcn4Δ*/*GCN4* strains on proline catabolic pathway metabolites, including arginine, ornithine, glutamate, and α-ketoglutarate. Utilization of urea and α-ketoglutarate was unaffected, as urea entered a separate catabolic pathway after arginine degradation (4). A mild defect was observed in glutamate utilization, likely due to *GDH2* being a target of *GCN4*-regulated amino acid control (GAAC) (data not shown). In contrast, arginine, ornithine, and proline utilization were defective in *gcn4Δ*, indicating that arginine and ornithine utilization are dependent on proline catabolism and that the defect lies specifically in the conversion of proline to glutamate via *PUT1* and *PUT2* (Fig. 3a-b). The absence of defects in α-ketoglutarate utilization supports a block at the level of proline catabolism.

**Figure 3.**
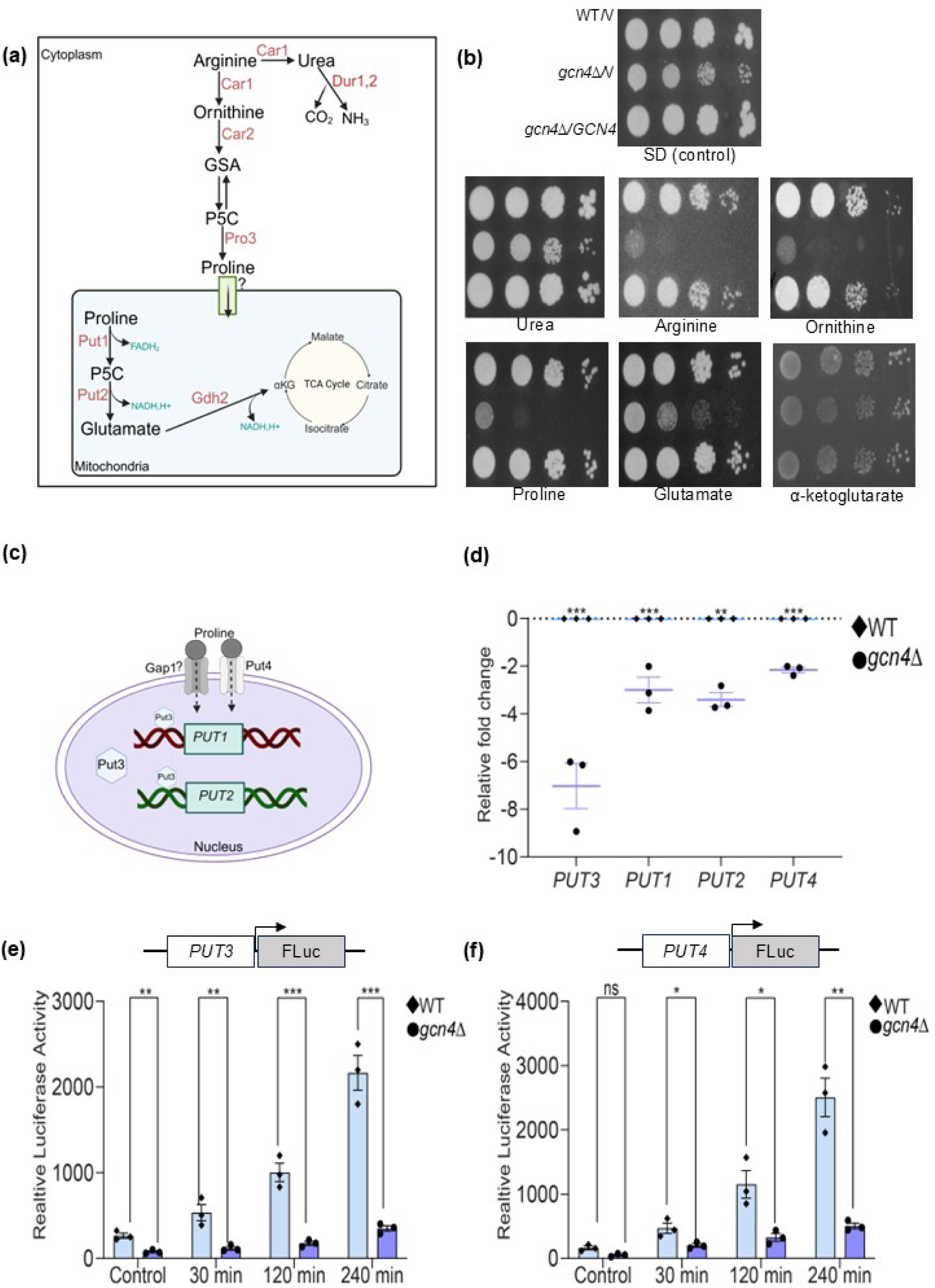
*GCN4* regulates the proline catabolism pathway via *PUT3* and *PUT4*. (a) Catabolism of arginine, ornithine, and proline. Arginine is converted to ornithine and urea by Car1, with urea subsequently processed by Dur1 and Dur2. Ornithine is converted to P5C by Car2, and Pro3 catalyzes the formation of proline from P5C. Proline is transported into mitochondria through an unidentified transporter, where it is sequentially oxidized to glutamate by Put1 and Put2. Glutamate is further converted to α-ketoglutarate by Gdh2, feeding into the Krebs cycle. Electrons generated as FADH₂ and NADH enter the mitochondrial electron transport chain. (b) The spot assay of WT, *gcn4Δ/V*, and *gcn4Δ/GCN4* grown on ammonium sulfate and on various metabolites provided as the sole nitrogen source that directly or indirectly influence the proline catabolism. (c) Proline uptake is mediated by permeases such as Gap1 (a general amino acid permease) and Put4 (a proline-specific permease). Proline entry into the cell maximally activates the transcription factor Put3, which upregulates the proline catabolism genes *PUT1* (proline dehydrogenase) and *PUT2* (1-pyrroline-5-carboxylate dehydrogenase), enabling the mitochondrial conversion of proline to glutamate. (d) Expression status of the *PUT* gene family in *gcn4Δ* compared to WT during proline utilization. WT and *gcn4Δ* strains were grown in minimal media with proline as the sole nitrogen source for 4 h. RNA was extracted at 240 min, and qRT-PCR was performed for *PUT3*, *PUT1*, *PUT2*, and *PUT4* using 5.8S rRNA gene and *UBC13* as internal controls. Mean ± SEM from three biological replicates is shown. (e-f) The promoter reporter constructs *PUT3*-Fluc and *PUT4-* Fluc were transformed into WT and *gcn4Δ* ura^-^ strains. Reporter activity for *PUT3* and *PUT4* expression was measured by a luciferase assay described previously, using ammonium sulfate (SD) as the untreated control and proline as the sole nitrogen source at 30 min, 120 min, and 240 min. The luciferase values were normalized to both the control reporter and total protein content, and the mean values from three independent biological replicates (n=3) were plotted. Statistical significance was determined using a two-tailed, unpaired Student’s *t*-test (*p<0.05, **p<0.01, ***p<0.001).

Furthermore, we analyzed the expression of proline catabolic pathway genes *PUT1*, *PUT2*, the proline transcriptional regulator *PUT3*, and the proline permease *PUT4* (Fig. 3c-d). Transcript analysis under proline induction revealed significant downregulation of all four genes in *gcn4Δ* strain relative to WT (Fig. 3d). The most significant downregulation was observed in *PUT3* (Fig. 3d). Notably, *PUT4* was also downregulated, suggesting *GCN4* regulates both proline import and catabolism.

To further elucidate the role of Gcn4 in regulating *PUT3* and *PUT4*, we cloned the promoters of *PUT3* and *PUT4* upstream of the luciferase construct. Under proline induction, both *PUT3* and *PUT4* promoter activity were higher in WT than in *gcn4*Δ at all time points (Fig. 3e-f). These results indicate that *GCN4* regulates both *PUT3* and *PUT4*. Significant promoter activity of both genes in cells grown on ammonium sulfate indicates that *PUT* genes may be largely independent of nitrogen catabolite repression in *C. glabrata*, as observed in *C. albicans* (4). Overall, *GCN4* orchestrates proline catabolism in *C. glabrata* by directly regulating the transcriptional regulator *PUT3* and the proline permease *PUT4*. These findings establish *GCN4* as a central regulator of proline catabolism, integrating proline uptake and mitochondrial catabolic pathways to support cellular nitrogen metabolism.

### *PUT3* drives proline catabolism while *PUT4* coordinates proline uptake and metabolic integration in *Candida glabrata*

*GCN4* plays a crucial role in proline catabolism, as it regulates the expression of both *PUT3* and *PUT4*. To determine whether the defect in the *gcn4Δ* mutant arises from impaired proline catabolism or transport, *PUT3* and *PUT4* were independently overexpressed under the constitutive *PDC1* promoter in the *gcn4*Δ background. Remarkably, constitutive expression of *PUT3* partially rescued the growth defect when *gcn4Δ* cells were grown on proline as the sole nitrogen source, whereas *PUT4* overexpression failed to restore the same (Fig. 4a–b). Growth curve analysis further supported these observations, showing that *PUT3*-overexpressing *gcn4Δ* cells exhibited significantly improved growth on proline compared to the parental *gcn4Δ* strain. In contrast, *gcn4Δ*/*PUT4* cells displayed altered growth compared to *gcn4Δ* mutant grown on proline (Fig. 4c-d). This intriguing phenotype prompted the hypothesis that *PUT4* overexpression in the absence of *GCN4* might enhance proline uptake beyond the catabolic capacity of cells, leading to intracellular proline accumulation and consequent toxicity. To validate this hypothesis, intracellular proline levels were quantified after 120 min of proline utilization, as 120 and 240 min yielded similar results. Consistent with impaired catabolism, *gcn4Δ* cells accumulated significantly more proline (24.33 ± 1.45 µg/ml) than WT (15.87 ± 0.70 µg/ml). Complementation of the *gcn4*Δ strain with *GCN4* restored catabolic activity and reduced intracellular proline content to 12.66 ± 1.76 µg/ml, which was lower than WT levels, suggesting the role of *GCN4* in maintaining proline homeostasis (Fig. 4e). Overexpression of *PUT3* in the *gcn4Δ* background (*gcn4Δ*/*PUT3*) similarly reduced intracellular proline content to 15 ± 1.72 µg/ml, comparable to proline accumulation in WT cells, and restored growth (Fig. 4a, c, & e). This suggests that *PUT3* overexpression can compensate for the loss of *GCN4*. Conversely, *gcn4Δ*/*PUT4* cells exhibited abnormally high intracellular proline levels (35.67 ± 4.98 µg/ml), far exceeding those noted in *gcn4*Δ cells, indicating enhanced uptake without corresponding catabolic processing (Fig. 4b, d, & e). Collectively, these findings demonstrate that *PUT4* primarily facilitates proline uptake, but in the absence of a functional catabolic system, increased transport exacerbates intracellular toxicity. They also suggest that *PUT4* is not the sole proline transporter in this fungus, as *gcn4Δ*/*PUT3* cells exhibit restored growth without the expression of *PUT4* in *the gcn4*Δ strain. In contrast, *PUT3*, a transcriptional activator of proline catabolic genes (*PUT1* and *PUT2*), plays a pivotal role in driving the degradation pathway essential for efficient proline utilization. Thus, while *PUT4* contributes to proline import and integration into the metabolic network, *PUT3***-**mediated regulation of catabolic enzymes is essential for channeling proline into central metabolism. Together, these findings establish that *GCN4* governs proline catabolism in *C. glabrata* primarily by transcriptionally activating *PUT3*, which orchestrates catabolic flux. In contrast, *PUT4* functions as a facilitator of proline uptake whose activity must be balanced with downstream degradation capacity to prevent toxic accumulation.

**Figure 4.**
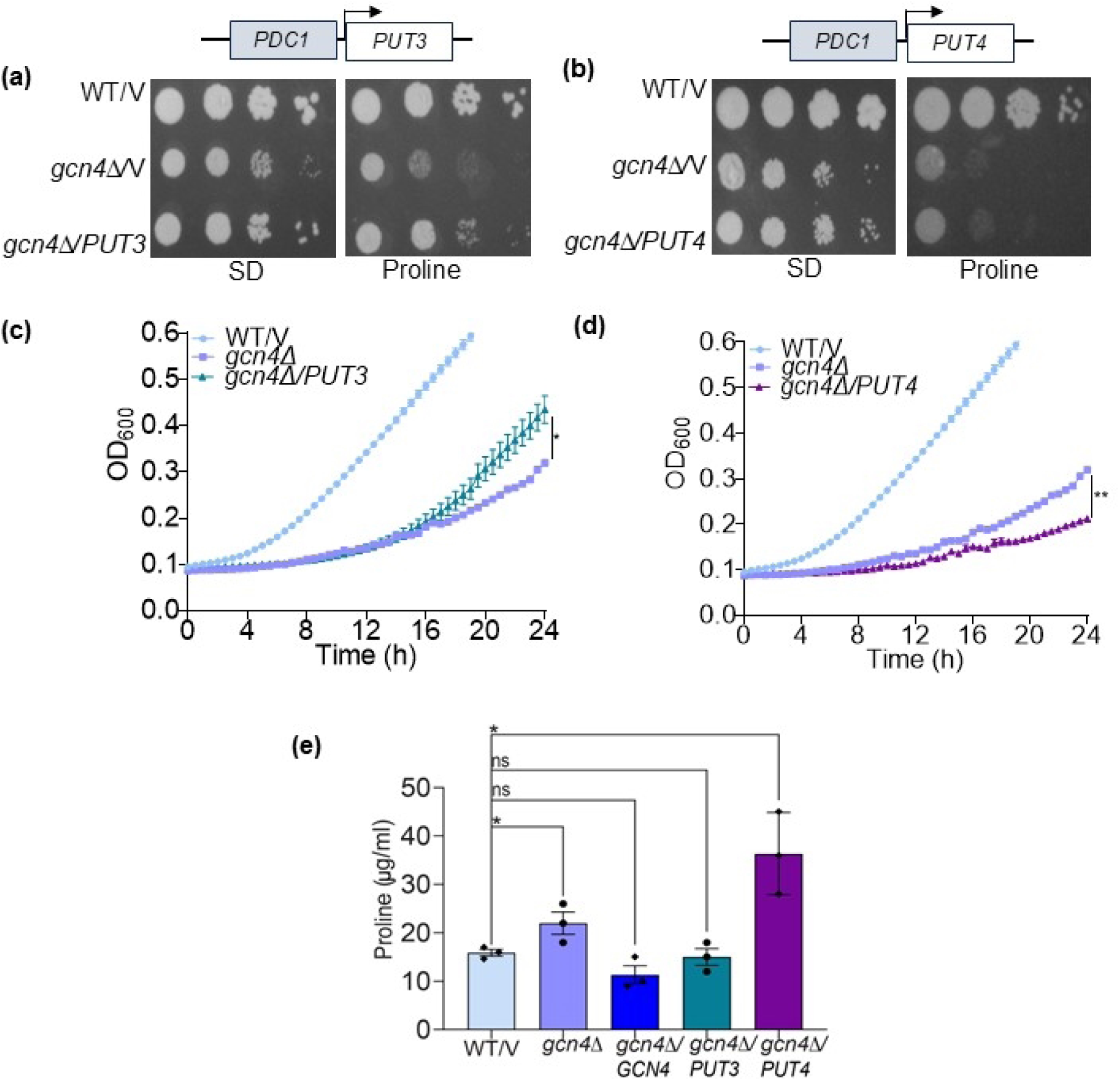
*GCN4-*regulated *PUT3* efficiently supports proline utilization compared with *PUT4*. (a-b) The genomic sequences (excluding UTRs) of *PUT3* and *PUT4* were cloned downstream of the constitutive PGK1 promoter and transformed into the *gcn4Δ* ura^-^ strains. The spot assays of WT/V, *gcn4Δ/*V, *gcn4Δ/PUT3*, and *gcn4Δ/PUT4* were performed on SD media containing ammonium sulfate and on minimal media with proline as the sole nitrogen source. (c-d) *Candida glabrata* WT/V, *gcn4Δ/*V, *gcn4Δ/PUT3*, and *gcn4Δ/PUT4* strains were grown in SD-Ura medium, harvested, and adjusted to an O.D. of 0.12 in a 96-well plate in minimal media containing proline as the sole nitrogen source. Growth was monitored every 30 min for 24 h using a Spectramax plate reader. The representative growth curves from three biological replicates (n=3) were plotted, and the AUC was calculated using Prism software. The mean ± S.E.M. was then plotted. (e) The colorimetric estimation of intracellular proline levels in WT/V, *gcn4Δ/*V, and *gcn4Δ/PUT3* and gcn4Δ/PUT4 strains following 2 h of proline induction is shown. Total proline was quantified using the ninhydrin assay and normalized to total protein. The proline concentration in each sample was plotted as µg/ml, and the data represent the mean of three biological replicates (n=3). Statistical significance was determined using a two-tailed, unpaired Student’s *t*-test (*p<0.05, **p<0.01, ***p<0.001).

### Gcn4 regulates proline catabolism by regulating *PUT3*

Put3 is necessary for efficient proline catabolism, as its expression complements the growth of *gcn4Δ* cells on proline. *Candida glabrata* Put3 shares approximately 46.5% sequence identity with its *S. cerevisiae* ortholog. It encodes a zinc cluster transcription factor that contains a characteristic Zn(II)₂-Cys₆ DNA-binding domain, a linker region, and a C-terminal dimerization motif (36). Both the Zn(II)₂-Cys₆ DNA-binding domain and the leucine zipper domain are highly conserved across fungal species (Fig. S3c). To investigate the GCN4 binding sites on *PUT3*, we analyzed its upstream regulatory region for potential GCN4-binding motifs. We identified a putative *GCN4*-binding site (TGASTCA) on the *PUT3* promoter, which closely resembles the canonical *S. cerevisiae GCN4* consensus sequence (37, 38). Further analysis revealed an eight-base motif “TGAGTCAG” located approximately 400 bp upstream of the *PUT3* ORF. Promoter sequences of *PUT3* homologs from non-pathogenic fungi like *S. cerevisiae* as well as pathogenic fungal spp., including *A. fumigatus*, *C. neoformans,* and several *Candida* spp. such as *C. albicans* and *C. auris*, were extracted and analyzed for the presence of Gcn4-like motifs. A highly similar motif was conserved across all the homologs, with A and C at -3 and -6 positions being most frequent (Fig. 5a-b). This strong conservation further suggested that Gcn4 binds to the *PUT3* promoter.

**Figure 5.**
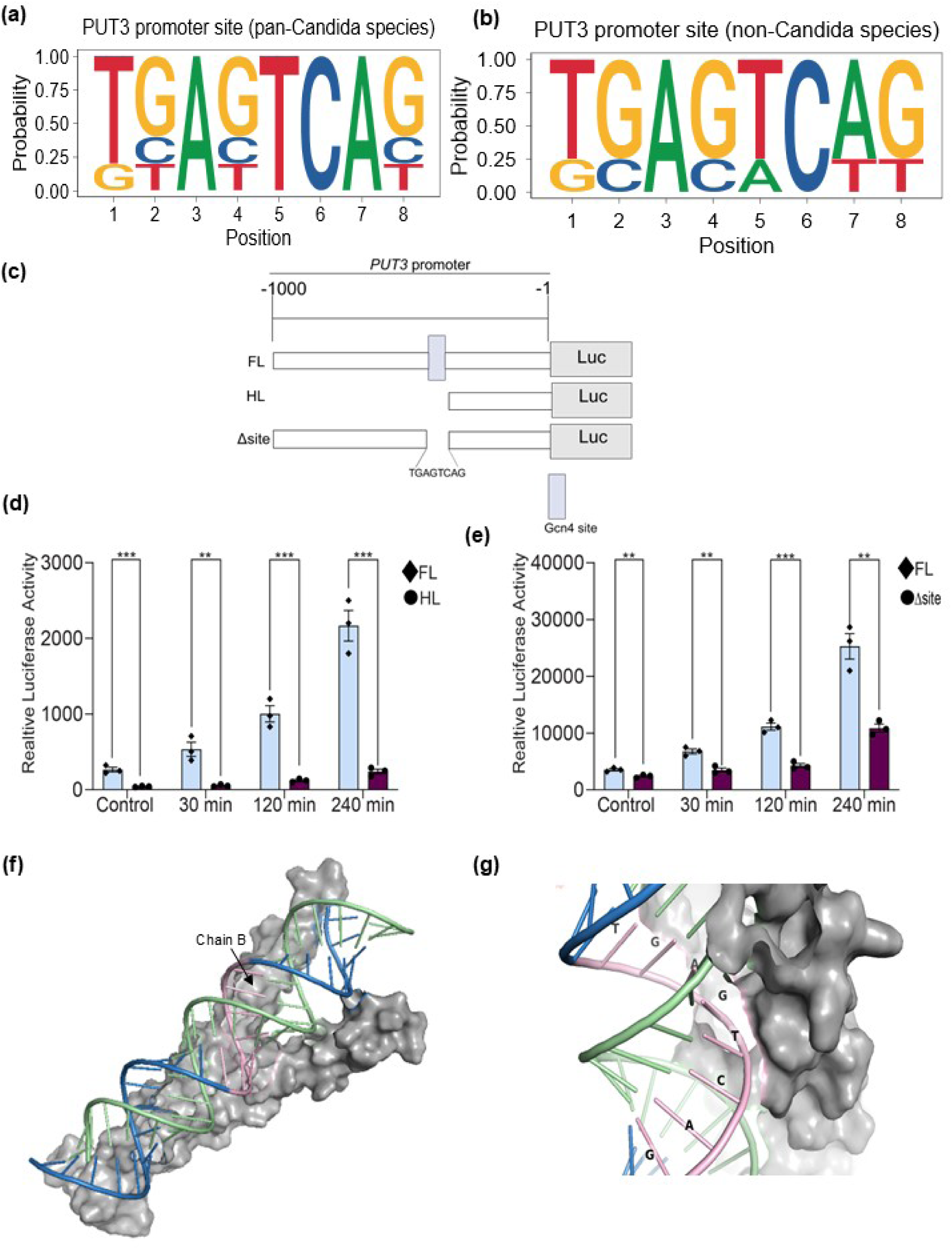
*GCN4* binds to the *PUT3* promoter. (a-b) Promoter analysis of *PUT3* orthologs in (a) pan-*Candida* species (*C. albicans*, *C. dubliniensis*, *C. glabrata*, *C. parapsilosis*, and *C. auris*) and (b) non-*Candida* species (*S. cerevisiae*, *S. pombe*, *A. fumigatus*, *C. neoformans*, and *K. lactis*). (c) Schematic representation of the *PUT3* promoter constructs. The full-length promoter (FL; -1000 to -1) was cloned upstream of the Fluc reporter gene. The half-length promoter construct (HL) lacks the region containing the predicted Gcn4 binding motif (GAGTCAG). A third construct (Δ site) carries the targeted deletion of the predicted Gcn4 binding site (highlighted in green) located at -606 to -601 bp. (d) Luciferase assay was performed using the *PUT3* FL promoter and the HL Fluc reporter constructs under control conditions (SD medium) and upon proline induction at 30, 120, and 240 min. (e) Luciferase assay of the *PUT3* FL promoter and the Δ site construct was similarly measured under control (SD medium) and proline as the sole nitrogen source at the same time points. All luciferase values were normalized to the control reporter and total protein content. Mean ± S.E.M. values from three independent biological replicates are shown. Statistical significance was determined using a two-tailed, unpaired Student’s *t*-test (*p<0.05, **p<0.01, ***p<0.001). (f) Molecular docking of Gcn4 leucine zipper and *PUT3* was performed using the HDOCK server, and the best model was selected based on high docking and confidence scores. (g) Binding interactions of the TGAGTCAG promoter motif of *PUT3* with the Gcn4 leucine zipper.

Promoter analysis also revealed binding sites of other transcription factors, including Gln3 and Stp2 (regulators of nitrogen), as well as stress-responsive factors such as Msn2 (a stress-responsive transcription factor), Mig1 (a regulator of glucose repression), and Skn7 (a regulator of oxidative stress), indicating a broader regulatory role of Put3 in stress responses. To examine the functional significance of the identified *GCN4*-binding site, we first generated a truncated *PUT3* promoter lacking approximately half of its upstream region, which included the putative binding site (Fig. 5c-d). The truncated promoter exhibited markedly reduced luciferase activity during proline utilization compared to the full-length construct, suggesting that this region contains crucial regulatory elements necessary for transcriptional activation of *PUT3* (Fig. 5d). However, since multiple cis-regulatory motifs may reside in this region, we next sought to specifically determine the contribution of the GCN4-binding site. Using overlap extension PCR, we deleted the specific eight-base pair motif (TGAGTCAG) corresponding to the predicted GCN4-binding site and cloned the modified promoter upstream of the luciferase reporter gene (Fig. 5c & e). The full-length promoter exhibited a significant increase in luciferase expression upon proline induction at different time points and showed 1.48-fold higher basal activity relative to the Δsite construct. This difference further increased on proline utilization, reaching 1.95-fold (30 min), 2.61-fold (120 min), and 2.33-fold (240 min) compared to the Δ site promoter (Fig. 5e). These results indicate that deletion of the GCN4-binding motif severely impairs proline-induced activation of *PUT3*, confirming that this element is critical for Gcn4-mediated transcriptional regulation during proline utilization.

To validate this interaction, we modeled docking between the Gcn4 leucine-zipper domain and the promoter region of *PUT3* DNA harboring the site. From the HDOCK docking results, we obtained the top 10 models of the Gcn4–*PUT3* complex. The model exhibited the highest docking and confidence scores; it contained a large number of interface residues contacting the TGAGTC motif of *PUT3*. The docking and confidence scores of the model were -253.87 and 0.89, respectively (Fig. 5f & g). A more negative docking score and a confidence score greater than 0.7 suggest that the Gcn4-PUT3 interaction is favorable. This model was subsequently used for molecular dynamics (MD) simulations (Fig. 5f & g). Additionally, a plot of the distances of the predicted GCN4-binding residues from the protein showed that G13 had the shortest distance, followed by T14, C15, and G11 (Fig. S2a).

Root mean square deviation (RMSD) analysis showed that the Gcn4–*PUT3* complex stabilized after approximately 200 ns and maintained a consistent conformation for the remainder of the 300-ns MD simulation. The TGAGTCAG motif displayed comparable stability (RMSD: 3.27 ± 1.41 Å), indicating no unusual conformational changes upon protein binding (Fig. S2b & c). Distance analysis of the final 100 ns identified two primary anchors and one secondary contact. ARG74 formed the strongest interaction with G127 (position 4), LYS85 showed comparable binding to T128 (position 5), and ARG66 exhibited moderate contact with G125 (position 2) (Fig. S2f). Comprehensive H-bond analysis identified 123 unique protein–DNA H-bonds with >20% occupancy. The binding residues showed distinct patterns: R66 formed 27 H bonds, R74 formed 3 H bonds, and K85 formed 9 H bonds to T128. The higher H bond occupancy for R66 (80.6%) compared to distance occupancy (13.5%) reflected the long, flexible arginine side chain forming H bonds at distances of 5–7 Å. Conversely, lower H bond occupancy of K85 (14.7%), despite high distance occupancy (57.3%), was attributed to lysine’s single donor versus arginine’s three donors, favoring electrostatic over H bond interactions (Fig. S2 g-h).

Moreover, root-mean-square fluctuation (RMSF) analysis revealed that protein binding substantially restricted DNA mobility. The TGAGTCAG motif exhibited an RMSF of 2.86 ± 0.62 Å, compared to 7.66 ± 3.25 Å for unbound DNA regions, representing approximately 2.7- fold reduction (Fig. S2d & e). Chain B maintained an average of 204 atomic contacts (<5 Å) with the TGAGTCAG motif across all frames, with 100% persistence, indicating no dissociation events during the 100-ns analysis window. Per-residue contact analysis identified R77 (96.0 contacts; 100% of frames), R74 (47.5 contacts; 99.8% of frames), and R66 (33.8 contacts;94.9% of frames) as the major contributors to the protein–DNA interface.

Overall, the simulation revealed asymmetric, non-canonical binding wherein chain B of Gcn4 leucine zipper engaged with promoter motif, primarily through N-terminal basic residues (R66, R74, and K85). The complex showed no dissociation over 300 ns, and the interface RMSD of 3.26 ± 1.38 Å indicated stable recognition of the *PUT3* motif. Collectively, these results demonstrate that the identified *GCN4*-binding site is both functionally and structurally relevant for binding of Gcn4 over *PUT3* promoter, highlighting the importance of *GCN4* in regulating proline catabolism.

### *PUT3* activates proline catabolism for the utilization of proline as a carbon and nitrogen source

Proline utilization involves the activation of transcriptional factor *GCN4*, which in turn regulates the another transcription factor *PUT3*. The deletion of *put3* impaired proline utilization as a sole nitrogen source (Fig. 6a & 4a), leading to intracellular proline accumulation (26 ± 2.3 µg/ml), which was restored upon complementation (16 ± 1.15 µg/ml) (Fig. 6b).

**Figure 6.**
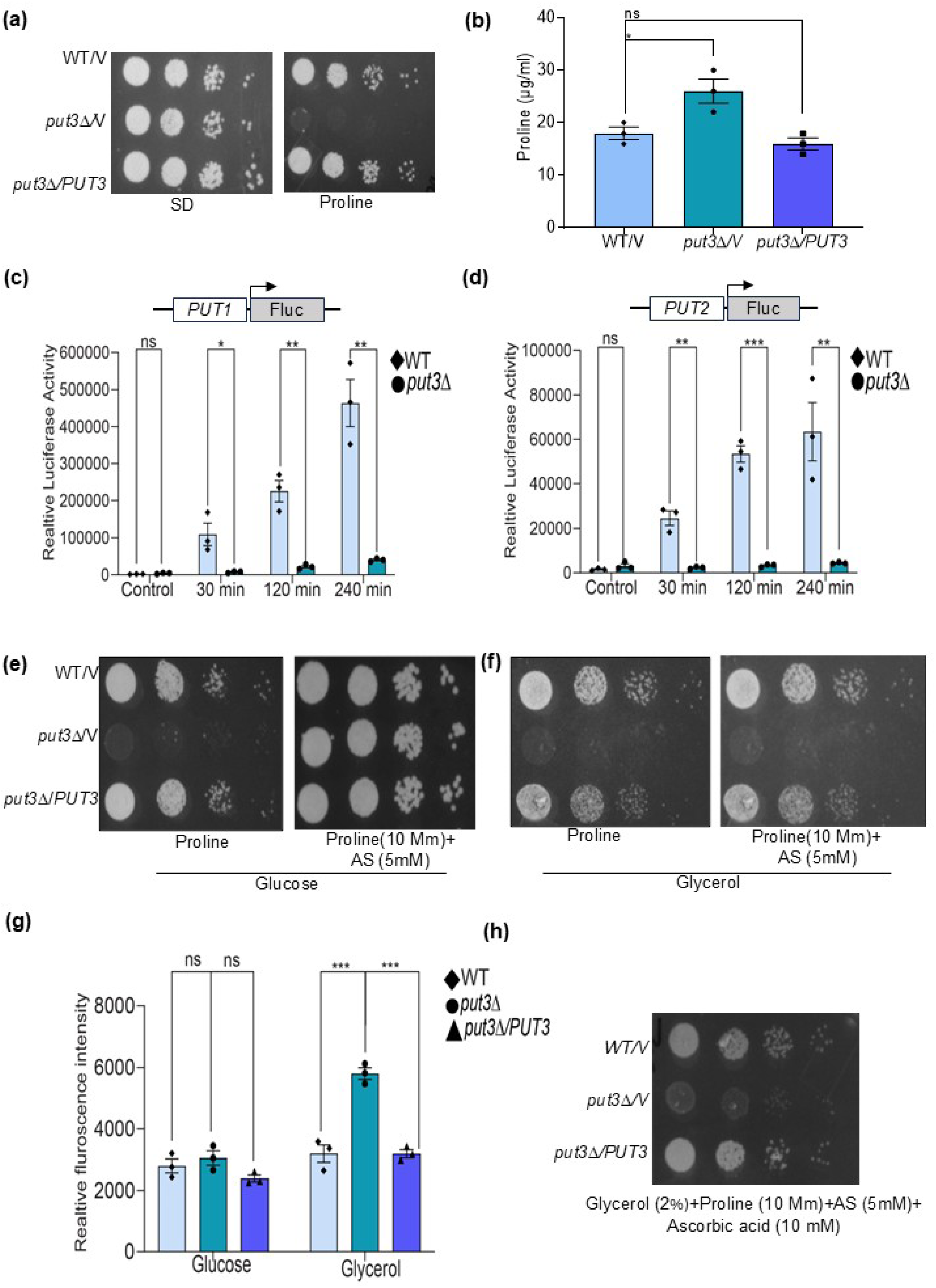
*PUT3* is required for proline utilization and regulates both the proline catabolic pathway and mitochondrial electron transport. (a) The result of spot assay of WT, *put3Δ,* and *put3Δ/PUT3* on SD media and on medium containing proline as the sole nitrogen source. (b) Colorimetric proline estimation of WT, *put3Δ*, and *put3Δ/PUT3* was performed as described in figure 4e. The total proline content (µg/ml) was normalized to total protein. Mean ± S.E.M. from three biological replicates is shown (n=3). (c-d) The promoter-reporter constructs *PUT1*-Fluc and *PUT2*-Fluc were transformed into WT and *put3Δ* ura^-^ strains. Reporter activity for *PUT1* and *PUT2* expression was measured through a luciferase assay using ammonium sulfate (SD) as the untreated control and proline as the sole nitrogen source at 30 min, 120 min, and 240 min. Luciferase values were normalized to the control reporter and total protein content. Mean values from three biological replicates were plotted (n=3). Statistical significance was determined using a two-tailed, unpaired Student’s *t*-test (*p<0.05, **p<0.01, ***p<0.001). (e) The results of spot assays of WT, *put3Δ*, and *put3Δ*/*PUT3* on media containing 2% glucose, with proline alone or proline supplemented with 5 mM ammonium sulfate, are shown. (f) The results of spot assays of the same strains on media containing 2% glycerol with proline alone or proline supplemented with 5 mM ammonium sulfate are shown. (g) ROS assessment in WT, *put3Δ*, and *put3Δ/PUT3* grown on 2% glucose or glycerol as the carbon source are shown. Log-phase cells grown in SD medium (2% glucose or glycerol) were shifted for 4 h to media containing proline as the sole nitrogen source and either glucose or glycerol as the carbon source. Cells were harvested, washed, adjusted to an O.D. of 1, and stained with 10 µM H2DCFDA for 45 min at 30°C. Fluorescence (excitation 485 nm; emission 520 nm) was measured using SpectraMax and normalized to total protein content (Bradford assay). Arbitrary fluorescence units (AFUs) normalized to protein content were plotted (mean ± S.E.M.; n = 3). Statistical significance was assessed using a two-tailed, unpaired Student’s *t*-test (*p<0.05, **p<0.01, ***p<0.001). (h) Results of spot assay of WT, *put3Δ*, and *put3Δ/PUT3* on medium containing proline + 5 mM ammonium sulfate as nitrogen sources and 2% glycerol as the carbon source, supplemented with 10 mM ascorbic acid are shown.

Activation of *PUT1* and *PUT2* by *PUT3* mediated the sequential conversion of proline to glutamate. To demonstrate the regulation of these genes by *PUT3*, the promoters of *PUT1* and *PUT2* were cloned upstream of a luciferase reporter, and their expression was monitored in WT and *put3Δ* strains. *PUT1*-*Luc* showed robust luciferase induction (55.1-, 113.5-, and 233.4-fold at 30, 120, and 240 min, respectively), and *PUT2*-Luc also exhibited substantial upregulation (15.8-, 34.3-, and 40.8-fold at corresponding time points). In the *put3Δ* background, luciferase activity was markedly reduced for both genes *PUT1* and *PUT2,* demonstrating that both were under strong *PUT3-*dependent transcriptional control during proline utilization (Fig. 6c-d).

We further analyzed the amino acid sequences of both Put1 and Put2 across *Candida* species and other fungal species as described in the methodology. The *C. glabrata* amino acid sequences from CAGL0M04499g (*PUT1*) and CAGL0D03982g (*PUT2*) genes showed 57.73% and 76.41% similarity to *S. cerevisiae* Put1 and Put2, respectively. For functional characterization, we generated deletion mutants of *PUT1* and *PUT2* using homologous recombination in *C. glabrata*. The resulting *put1*Δ and *put2Δ* strains exhibited impaired growth on proline as the sole nitrogen source, confirming their role in its catabolism (Fig. S3a-b & S4b-e). Given that proline catabolism governs ornithine utilization, we assessed the regulation of *PUT1*, *PUT2*, and *PUT3* when ornithine was utilized as the sole nitrogen source. During ornithine utilization, *PUT1* was strongly induced (approximately 12-fold at 0–30 min), with sustained expression at later time points. *PUT2* also showed consistent upregulation (4.36- and 3.13-fold) over time. Interestingly, *PUT3* was also upregulated (approximately 2-fold) (Fig. S4f-h). In *put3Δ* strain, promoter activities of *PUT1* and *PUT2* significantly decreased in the presence of ornithine (Fig. S4i-j). Further growth defects were observed in *put3Δ* and *put2Δ* strains upon ornithine utilization (Fig. S4 l-m). However, *put1*Δ showed a less severe phenotype, possibly due to ornithine-to-proline conversion via ornithine transaminase, bypassing the *PUT1* requirement (Fig. S4 k) (39). Additionally, *PUT* mutants did not exhibit any defects in the utilization of urea, glutamate, α-ketoglutarate, or citrulline (data not shown).

Together, these results establish *PUT3* as the central coordinator of *PUT1* and *PUT2*. The metabolic link between proline and ornithine becomes especially clear as proline-derived glutamate contributes to ornithine biosynthesis via ornithine aminotransferase, highlighting a tightly integrated and interdependent pathway network (4) .

The proline catabolic pathway is closely linked to mitochondrial function due to the localization of its enzymes and the generation of reducing equivalents (FADH₂ and NADH), which feed into the electron transport chain (ETC) to drive oxidative phosphorylation and ATP synthesis (4, 24). *C. glabrata*, similar to *S. cerevisiae*, is a Crabtree-positive yeast that primarily relies on glycolysis for ATP generation when glucose is the main carbon source (27, 40). Under such conditions, mitochondrial activity is repressed, reducing the dependence of the organism on oxidative phosphorylation and bypassing the need for energy generation through the proline pathway.

Consistently, *put1Δ*, *put2Δ*, and *put3Δ* mutants exhibited growth similar to WT on glucose-containing media supplemented with proline and ammonium sulfate (5 mM) (Fig. 6e & S5a-b). This suggested that under fermentative conditions, the presence of ammonium compensated for the nitrogen contribution from proline.

However, under non-fermentative conditions (established using 2% or 4% glycerol as the carbon source), glycolysis was repressed, shifting the energy dependence toward mitochondrial respiration. Under these conditions, *put3Δ* displayed significant growth inhibition on media containing proline, which persisted even with the addition of ammonium sulfate (Fig. 6f). The *put1Δ* and *put2Δ* mutants also showed growth defects under the same conditions, albeit to a slightly lesser extent than *put3*Δ (Fig. S5c-d). These findings highlight the requirement of *PUT3*-mediated proline catabolism for efficient ETC function and ATP generation when glycolytic ATP production is limited.

Furthermore, *put3Δ*, when compared to WT and the complemented strain, exhibited approximately 80% higher reactive oxygen species (ROS) when grown on media containing glycerol and proline, whereas, under all other conditions, ROS levels were more or less similar (Fig. 6g). The increased ROS levels were likely due to the absence of Complex I in *C. glabrata*, which may compound the effects of defective proline catabolism on mitochondrial function and redox balance (41). Supplementation with antioxidants, such as ascorbic acid, did not rescue the growth defect in *put3*Δ (Fig. 6h). This suggests that oxidative stress alone is insufficient to explain this phenotype and instead reflects the toxic accumulation of P5C resulting from inefficient *PUT2* activity. This suggests a vital role for *PUT2*, regulated by *PUT3*, in P5C detoxification and ROS homeostasis. These findings collectively demonstrate that, despite its Crabtree-positive nature and lack of Complex I, *C. glabrata* relies on functional proline catabolism for efficient energy production and ROS management under respiratory growth conditions.

### Proline catabolism is essential for *Candida glabrata* virulence and pathogenicity

*Candida glabrata* can survive and replicate within host macrophages, a trait attributed to its efficient stress adaptation mechanisms, with Gcn2 and Gcn4 playing prominent roles (17). To further investigate whether *PUT3*-regulated proline metabolism contributes to this survival, we infected PMA-treated THP-1 macrophages with phototrophic WT and *put3*Δ strains of *C. glabrata* at a multiplicity of infection (MOI) of 1:10. Following a 2-h incubation to allow phagocytosis, intracellular yeast cells were collected at time points consistent with a previous study, capturing both early and later stages of infection (Fig. 7a) (17).

**Figure 7.**
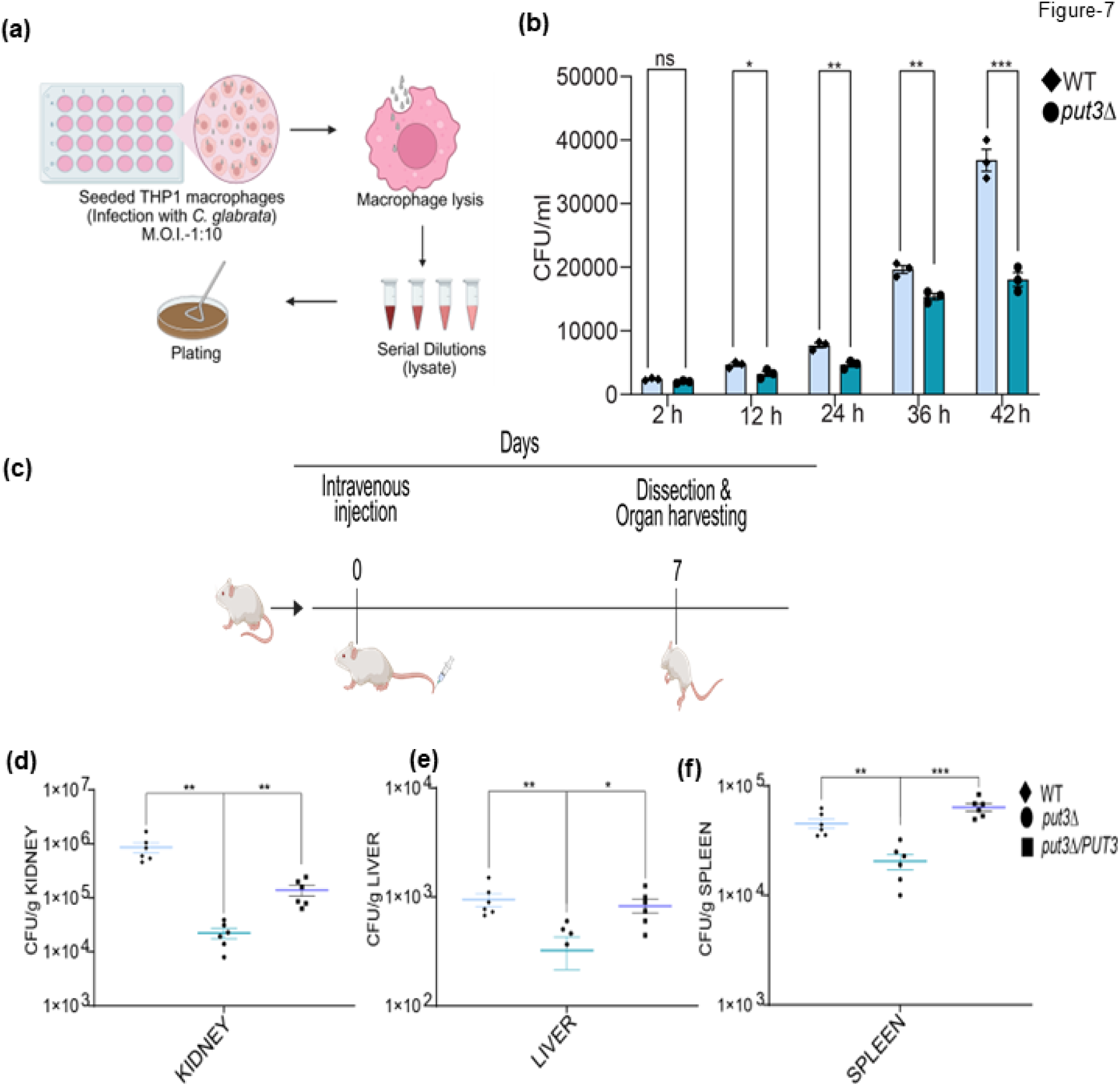
*PUT3* governs macrophage survival and host virulence. (a) Schematic of the macrophage infection assay. Macrophages were seeded and infected with yeast cells; after infection, macrophages were lysed, and internalized fungal cells were serially diluted and plated on YPD to determine CFUs. (b) Phototrophic *C. glabrata* Bg2 WT and *put3Δ* strains were infected into PMA-treated THP-1 macrophages at an MOI of 1:10 for 2 h. After infection, extracellular yeast cells were removed, and at the indicated time points, intracellular yeast cells were recovered by lysing the macrophages, serially diluted, and plated on YPD. Plates were incubated at 30°C for 2–3 d. Infection experiments were performed in triplicate for each strain, and CFU/ml (mean ± S.E.M.) values were calculated from three independent experiments (n=3). Statistical significance was determined using a two-tailed, unpaired Student’s *t*-test. (c) Schematic of the disseminated candidiasis murine model. Female BALB/c mice were intravenously injected with 1×10⁸ fungal cells and monitored for 7 d, after which target organs were collected for fungal burden analysis. (d-f) Virulence assessment of *C. glabrata* Bg2 WT and *put3Δ* strains, along with *put3Δ/PUT3*- complemented strains, using the disseminated candidiasis model. Female BALB/c mice were intravenously infected with 1×10⁸ cells for 7 d. Mice (n = 6 per strain across two independent experiments) were sacrificed, and the kidneys (d), liver (e), and spleen (f) were aseptically removed, weighed, homogenized, and plated on YPD supplemented with penicillin and streptomycin. Plates were incubated at 30°C for 3–4 d, after which CFUs were counted and expressed as CFU/g of tissue. Data represent six individual mice per strain (mean ± S.E.M.). Statistical significance was determined using a two-tailed, unpaired Student’s *t*-test (*p<0.05, **p<0.01, ***p<0.001).

Despite similar phagocytosis rates between WT and *put3Δ*, the CFU/mL of the *put3Δ* was consistently reduced at all time points (Fig. 7b). As early as 12 h post-infection, the *put3Δ* showed a 31% reduction in CFUs, which increased to approximately 51% by 42 h compared to WT (Fig. S6a). In terms of replication, WT cells showed an increase from 2.5-fold at 12 h to 15-fold at 42 h, while *put3Δ* showed a more modest increase from 1.5- to 9-fold over the same period (Fig. S6b). Notably, the fold replication of *put3Δ* plateaued between 36 h (8-fold) and 42 h (9-fold), whereas WT replication continued to rise sharply from 8- to 15-fold during that time period (Fig. S6b). This modest increase in mutant CFUs and the pronounced difference in fold replication at 42 h highlight the critical role of alternative nitrogen source utilization, particularly proline metabolism regulated by *PUT3*, in sustaining long-term intracellular survival and replication of *C. glabrata* within the nutrient-limited environment of host phagosomes.

To assess the contribution of *PUT3* to *C. glabrata* virulence, we employed the intravenous murine candidiasis model, which enables systemic fungal dissemination via direct inoculation into the bloodstream (Fig. 7c). As anticipated in this model, kidney, the principal target organ, exhibited the highest fungal burden in mice infected with the WT strain. Given the availability of proline in kidneys, this organ serves as a biologically relevant site for assessing the role of *PUT3* in host–pathogen interactions (7). Thus, female BALB/c mice were injected via the tail vein with 1 × 10⁸ CFUs of WT, *put3Δ*, or complemented (*put3Δ*/*PUT3*) strains. After seven days, animals were euthanized, and fungal burdens in the kidney, liver, and spleen were quantified by plating serial dilutions of homogenized tissue. Uninfected animals served as negative controls.

The *put3*Δ-infected mice showed a significant reduction in fungal burden in all examined organs relative to the WT-infected mice. In the kidneys, the fungal load decreased by approximately 36-fold, while reductions of 21-fold and 3-fold were observed in the liver and spleen, respectively. Complementation restored fungal colonization, with approximately 5- and 2.6-fold increases in the kidney and spleen, respectively, relative to the mutant. Liver colonization also increased in the complemented strain (approximately 5-fold) compared to *put3*Δ (Fig. 7d-f). *PUT3* has also been found to be upregulated in in vivo transcriptomic datasets (42), supporting its functional relevance during infection. These findings demonstrate that *PUT3* is critical in systemic virulence, likely through its involvement in mitochondrial proline catabolism, within the organ microenvironment.

Given that *PUT3* is essential for both in-vitro and in-vivo survival in *C. glabrata,* and it lacks a human homologue while remaining highly conserved within the fungal kingdom, it represents a promising fungal-specific therapeutic target. To explore the therapeutic potential of Put3 from *C. glabrata,* its binding sites were predicted using PrankWeb and DoGSiteScore to identify putative druggable pockets. PrankWeb identified Pocket 1 as the most probable (0.754) and the highest-scoring pocket (18.53), while DoGSiteScore identified Pocket P_1 (DrugScores > 0.8 and SimpleScores > 0.6) as highly druggable (Fig. S7 a-b). Collectively, the analyses from both the software suggest that ligands with a predominantly hydrophobic scaffold, complemented by a strategically positioned polar or basic functional group, may enable directional hydrogen bonding and electrostatic interactions within the Put3 binding pockets, likely enhancing binding affinity and specificity.

## DISCUSSION

Translational control enables cells to rapidly reprogram gene expression in response to nutrient and environmental stress. In this study, we identified a previously unknown regulatory axis linked to proline catabolism in *Candida glabrata*. Although proline catabolism has been thoroughly investigated across various systems, limited information is available about the specific signaling pathways and regulatory mechanisms that govern it. In this regard, we provided the first systematic and mechanistic evidence that a nutrient-responsive transcription factor is activated by translational control to coordinate gene expression and metabolism. We show that proline utilization represses global protein synthesis by phosphorylating eIF2, thereby decreasing ternary complex formation. This translational repression triggers Gcn4 activation which, in turn, selectively induces the expression of the transcription factor Put3 and Put4. Put3 then regulates downstream catabolic enzyme-coding genes of the proline pathway, including *PUT1* and *PUT2*, for the effective conversion of proline to glutamate (Fig. 8, upper left). This mechanism is even more crucial when proline is the sole nitrogen source, because fungal growth is markedly impaired with intracellular proline accumulation due to severe defects in proline catabolism caused by the deletion of either *PUT3* or its upstream regulator *GCN4*. In embryonic stem cells, proline has been reported to modulate the Gcn2-ATF4 pathway, thereby activating the amino-acid starvation response (AAR) pathway and regulates proline biosynthesis. However, the conclusion of this studies indicate that embryonic stem cells cultured in complete medium experience an intrinsic L-proline shortage that activates the AAR pathway, limit proliferation and suppresses the embryonic stem to mesenchymal transition, with this stress being alleviated by exoneneous L-proline supplementation.. Collectively, this finding primarily define proline as a starvation-associated metabolic signal that triggers the classical AAR pathway to activate the proline biosynthesis pathway to constrain cellular growth. However, the regulation of proline catabolism, its gene expression dynamics, and the downstream regulatory networks linking proline availability to activation of catabolic pathways remain largely unexplored.(43). In contrast our findings highlight proline utilization as a hierarchical regulatory framework in which *PUT3* functions as a specific transcriptional effector triggering the proline catabolic machinery. Here, we present the first integrated analysis of proline metabolism, defining the regulatory network that coordinates proline transport, assimilation, and downstream catabolic processes.. are (Fig. 8 upper).

**Figure 8.**
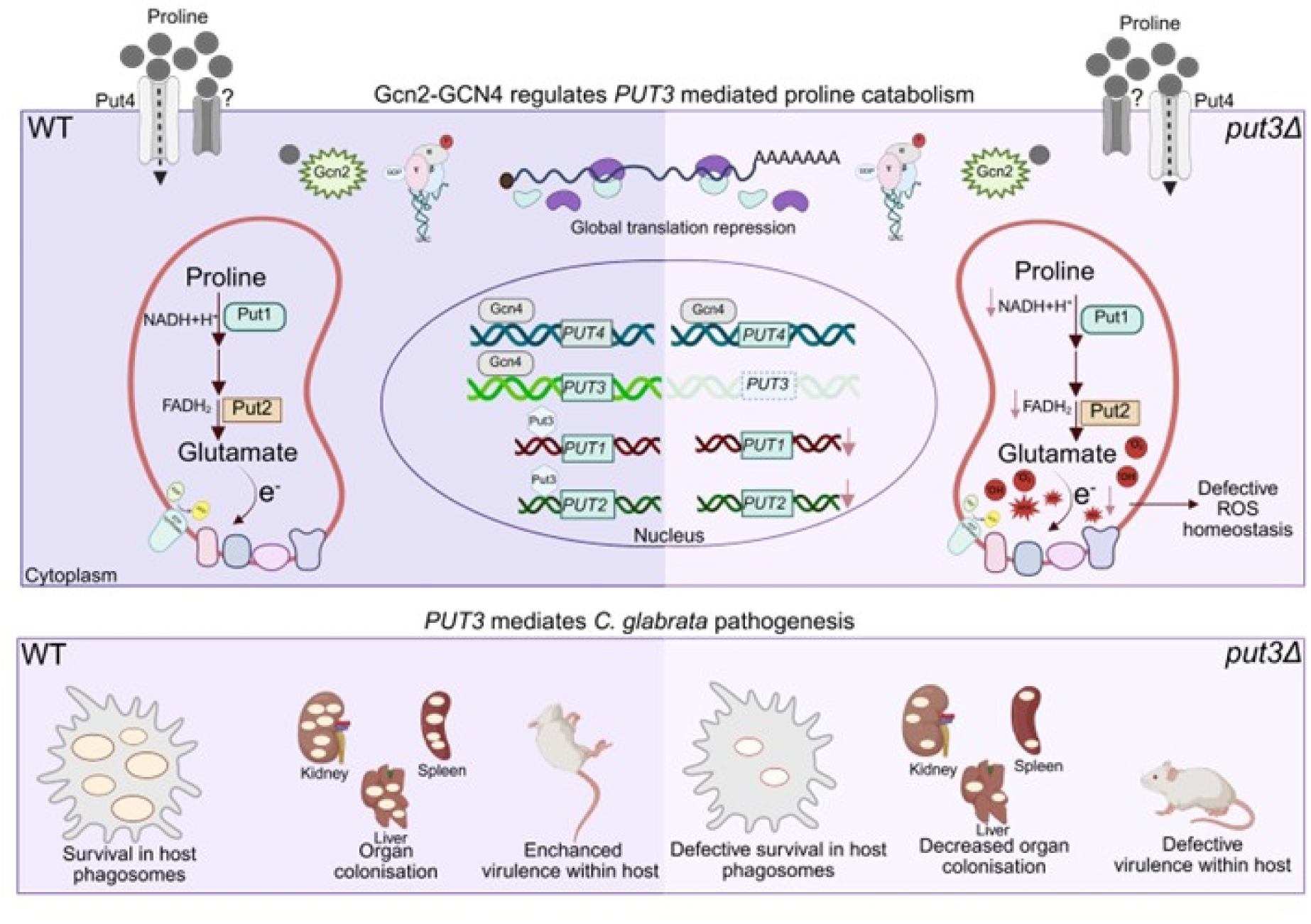
Gcn2–Gcn4–Put3 regulatory relay drives proline catabolism, ROS homeostasis, and virulence in *Candida glabrata*. (Top) Upon proline utilization, Gcn2 kinase is activated and phosphorylates eIF2α, leading to global translation repression and enabling the preferential translation of the transcription factor Gcn4. Gcn4 induces the expression of the proline transporter *PUT4* and the regulator *PUT3.* In WT *C. glabrata*, Put3 transcriptionally activates the proline catabolic genes *PUT1* and *PUT2*. Sequential oxidation of proline by Put1 (NADH-generating) and Put2 (FADH₂-generating) produces glutamate and donates electrons to the mitochondrial respiratory chain, enabling efficient proline catabolism and controlled ROS production. Thus, Gcn2-mediated translational control, together with Gcn4–Put3 transcriptional regulation, sustains efficient proline catabolism. Loss of PUT3 prevents the induction of proline catabolic genes downstream of the Gcn2–Gcn4 axis, resulting in impaired proline utilization, mitochondrial electron leakage, and increased ROS accumulation. (Bottom) These metabolic defects reduce fitness in host environments, as WT cells exhibit efficient survival within macrophage phagosomes, robust colonization of the liver, kidney, and spleen, and high virulence in a murine systemic infection model. In contrast, *put3Δ* cells display defective phagosome survival, reduced organ colonization, and attenuated virulence. Together, the model highlights a nutrient-sensing-to-translation control and transcriptional regulation cascade that safeguards fungal metabolism, ROS balance, and virulence within the host, also highlighting the central importance of the proline catabolic pathway in host adaptation. Figure created with biorender.com (https://www.biorender.com/).

Functionally, Put3 regulate the proline catabolism by regulating the expression of Put1 and Put2, mitochondrial enzymes involved in the catabolism of proline to glutamate, as observed in *S. cerevisiae* and *C. albicans* (44, 45). In our study, we found that *PUT1* and *PUT2,* in addition to proline, are also regulated during ornithine utilisation in a *PUT3*-dependent manner, and all *put* mutants exhibit growth defects upon ornithine utilisation, consistent with the fact that ornithine metabolism is solely dependent on proline catabolism (4). In this study, the *gcn4Δ* strain exhibited severe growth defects on arginine and ornithine, highlighting Gcn4’s broader role in coordinating these interconnected pathways. Along with the Gcn4-*PUT3*/*PUT4* regulatory cascade, this study also outlines the functional significance of proline uptake versus catabolism. *PUT4* facilitates proline transport, hazardous intracellular proline buildup results from its overexpression and in *gcn4Δ* in the absence of a functional catabolic pathway, demonstrating that absorption alone is insufficient for metabolic adaptation. Conversely, proline consumption is restored by constitutive expression of *PUT3* in the *gcn4Δ*, indicating that the primary limiting factor in this system is transcriptional activation of catabolic genes. It demonstrates the distinct yet complementary roles of transport and catabolic regulation, providing a comprehensive framework for understanding proline metabolism in fungal pathogens.

*PUT4* is not the sole proline transporter, as the *gcn4Δ* strain accumulates sufficient intracellular proline that can be further utilized upon *PUT3* overexpression. This indicates the presence of alternative proline transporters that contribute to proline uptake. Similar redundancy in proline transport systems has also been reported in other organisms, such as *C. albicans* (46). In the present study, overexpression of *PUT3* in the *gcn4*Δ background complemented the strain and reduced intracellular proline accumulation, further emphasizing the central role of catabolic regulation. Strikingly, *PUT3* overexpression rescued both the growth defects and proline accumulation observed in *gcn4*Δ, which were caused by alterations in both proline transport and catabolism. However, the growth of the *gcn4Δ*/*PUT3* strain could not fully mirror WT levels, likely because *PUT4* induction was absent, which limited proline import. Alternatively, as Gcn4 is a master regulator of stress-responsive pathways, other downstream pathways may also be compromised in the *gcn4Δ* strain, resulting in only partial rescue. In contrast, overexpression of *PUT4* alone led to excessive proline accumulation and further impaired growth, even more than in the *gcn4Δ* strain, indicating that proline toxicity arises when uptake exceeds catabolic capacity. These findings underscore the necessity of a balanced coordination between nutrient import via *PUT4* and catabolic processing mediated by *PUT3* to ensure efficient proline utilization and cellular homeostasis.

This critical metabolic fine-tuning may be more beneficial for *C. glabrata*, which lacks hyphal formation, a known virulence trait, and relies on efficient stress adaptation to survive in nutrient-limited conditions. The observations noted in this study highlight the role of *PUT3* as a key downstream effector of Gcn4-mediated catabolic regulation. Thus, based on the findings, we further investigated the physiological and pathogenic relevance of *PUT3* in *C. glabrata*.

Transcriptomic analysis of WT and *gcn4*Δ revealed *PUT3* as a target of Gcn4 (17). In this study, we observed that *gcn4Δ* exhibited defective proline utilization, and Gcn4 was regulated by Gcn2 kinase. Additionally, the *GCN4*-Luc reporter was upregulated upon exposure to proline. Gcn2 may thus act as a proline sensor, assisting in the activation of key regulators required for efficient proline utilization. *GCN4* regulates *PUT3* by binding to its promoter through a conserved palindromic motif, TGA(C/G)TCAG (37, 38). This conserved sequence has been detected across a variety of non-pathogenic and pathogenic fungal species, highlighting that Gcn4 broadly regulates proline catabolism via Put3. This interaction was verified by luciferase reporter assays, as deletion of this motif significantly reduced *PUT3* promoter activity during proline utilization, establishing a direct regulatory link. Although residual expression was observed, possibly mediated by regulators such as Mig1, Gat1 (carbon/nitrogen signaling), and Msn2/Skn7 (oxidative stress), *PUT3* serves as a convergence point for both nutrient- and stress-responsive signaling pathways. Additionally, molecular docking analysis showed that the TGAGTC motif on promoter of PUT3 interact with Gcn4. Furthermore, molecular dynamics simulation analysis revealed the structural stability of the complex, and distance analysis showed that chain B of Gcn4, specifically residues R66, R74, and K85, was primarily responsible for binding to the *PUT3* promoter motif. The combined stability of the interface, as measured by docking and MD simulations, demonstrated that Gcn4 engages the TGAGTCAG motif in a stable, sequence-specific manner, validating it as a functional binding site within the *PUT3* promoter.

Similar to *C. albicans*, proline catabolism plays a role in mitochondrial ATP synthesis in *C. glabrata* (4, 7). In conditions where glucose is inhibited, its significance becomes clear. Ammonium sulfate supplementation in proline and glycerol media did not reverse the growth abnormalities of *put* mutants, although *C. glabrata* is a Crabtree-positive yeast. This suggests that proline catabolism plays a crucial role in energy metabolism. Furthermore, *PUT3* deletion resulted in elevated ROS levels under similar non-fermentative conditions, and antioxidants, such as ascorbic acid, were unable to counteract this, suggesting an underlying redox imbalance likely caused by P5C buildup (Fig. 8, upper and lower). Thus, *PUT3* appears to integrate nitrogen and carbon metabolism, regulating mitochondrial activity and facilitating energy production when glucose, the preferred carbon source, is unavailable.

The present study highlights both the conserved and divergent aspects of proline regulation among species. While *C. glabrata* utilizes proline effectively, like *C. albicans*, *PUT3* plays a more indispensable role in the metabolism and pathogenicity of *C. glabrata*. In *C. albicans*, growth is maintained on glycerol–proline medium supplemented with 5 mM ammonium sulfate, whereas the deletion of *PUT3* results in only partial growth abnormalities on proline– dextrose media (7). In contrast, *C. glabrata put3Δ* mutants show a total growth deficiency under both conditions.

The glucose–proline-insensitive phenotype of the *put3* mutant in *C. albicans* suggests that *PUT1* and *PUT2* operate independently in glucose-based media (7). However, a significant decrease in promoter activity, especially of the *PUT1* promoter, even when dextrose was the carbon source, indicated that *C. glabrata PUT1* and *PUT2* are highly dependent on *PUT3*. The role of *PUT3* in *C. glabrata* virulence further emphasizes its functional relevance. *PUT3* deletion in *C. albicans* did not affect mouse survival and restored CFUs comparable to WT. Additionally, *PUT2* contributes to virulence in both *C. albicans* and *C. neoformans*, most likely due to toxic P5C accumulation (7, 23). In contrast, *C. glabrata* shows a significant reduction in virulence when *PUT3* is deleted (Fig. 8, lower right). All of these findings point to *PUT1* and *PUT2* being more tightly regulated by *PUT3* in *C. glabrata*, which is in line with the regulatory cascade in *S. cerevisiae* (44).

*PUT3* phenocopies *PUT2* in both metabolic and virulence contexts, despite lacking functional similarity. A comprehensive targeted study of *PUT3* is therefore warranted. Due to variations in host colonization tactics, immunological interactions, and nutrient intake, the divergence in *PUT3* function between *C. glabrata* and *C. albicans* most likely represents an evolutionary rewiring of metabolic regulation (47). The regulatory circuit from *S. cerevisiae*, where transcriptional regulation by *PUT3* regulates mitochondrial proline catabolism, seems to be retained in *C. glabrata*. Crucially, virulence and metabolism *in C. glabrata* are linked via *Gcn4* and *PUT3*. The deletion mutants of both genes exhibit abnormalities in the colonization of mouse organs and macrophage survival, underscoring proline metabolism as a crucial virulence factor (Fig. 8, lower). This process has a particularly pertinent setting in the kidney, an organ rich in proline (48). Hence, *C. glabrata* relies on metabolic flexibility and nutrient adaptation for survival in the host environment. *PUT3* thus emerges as a central node that connects amino acid sensing, energy metabolism, redox balance, and virulence, thereby assisting *C. glabrata* in host persistence.

The rise in antifungal resistance, especially in non-*C. albicans* species like *C. glabrata*, whose resistance is increasing to common drugs like azoles and echinocandins (25, 49), is a serious concern. An alternative antifungal strategy could involve targeting virulence factors that aid pathogen invasion and persistence, such as hyphal formation, adhesion, or biofilm formation. While these are classic targets, equally potent but less explored options include metabolic regulators and transcription factors that control macro- or micronutrient pathways. Targeting these regulators could reduce selective pressure, minimize toxicity, and expand the antifungal drug arsenal for these species (20, 50). In this study, we have shown that *PUT3* regulates both central metabolism and virulence traits, is fungal-specific, and belongs to the binuclear zinc cluster family [Zn(II)₂-Cys₆]. These features make Put3 an attractive therapeutic target, a notion supported by the bioinformatic predictions indicating the presence of druggable binding pockets.

Compared to genes like *PUT1* or *PUT2*, targeting a central regulator like *PUT3* may offer broader efficacy by disrupting multiple downstream processes simultaneously. Additionally, *PUT3*, which is critical for fungal survival yet absent in humans, could minimize off-target effects. The findings of the present study also position *PUT3* as a fungal-specific transcriptional node with both metabolic and virulence roles. Additionally, the conservation of Put3, along with the Gcn4–Put3 regulatory network, across fungal species suggests that this mechanism may be a general strategy by which eukaryotic microbes adapt to nutrient stress.

Though the present study identifies the Gcn2–Gcn4–Put3 cascade as a key link between translation, proline metabolism, and virulence, further investigations are necessary to examine how other nutrient-sensing pathways, such as TOR and Snf1, interact with this circuit under various stress conditions. Moreover, ChIP-sequencing studies need to be conducted for defining the Put3 regulon, along with proteomic and metabolomic analyses to connect gene regulation to metabolic flux. Together, these investigations will further clarify how fungal pathogens coordinate translational reprogramming with metabolic adaptation for survival in the host.

## MATERIALS AND METHODS

### Strains, growth, and media conditions

*Candida glabrata* strains used in this study are listed in Supplementary Table S1. All strains are derivatives of the BG2 background. The cultures were maintained either in rich YPD (2% peptone, 2% dextrose, and 1% yeast extract) or minimal SD media (0.5% ammonium sulfate, 0.17% yeast nitrogen base, and 2% dextrose). For Bg14 (auxotrophic), SD with 0.008% uracil was utilized at 30°C. Amino acid induction/utilisation assays used SD medium without ammonium sulfate supplemented with 10 mM amino acids, except arginine, whose concentration was 4 mM or 10 mM. Unless otherwise specified, dextrose was used as the energy source. For specific assays, 5 mM ammonium sulfate and 2% glycerol were used. For plasmid propagation, we used the *Escherichia coli* strains DH5α and DH10 B, which were maintained in Luria–Bertani medium at 37°C. The spot assays and growth analyses were performed using cultures grown overnight in liquid YPD/SD media at 200 rpm. For spot assay cultures, the cells were pelleted and washed twice, and then set to an OD_600_ of 1. Four 10-fold serial dilutions were prepared, and 5 µL of each was spotted onto the respective media plates. For growth curve analysis, the overnight cultures were diluted in fresh SD media and grown to mid-log phase (OD₆₀₀ ≈ 0.7–1). Cells were pelleted, washed, and adjusted to an OD_600_ of 0.12 in 200 µl of SD medium or SD medium lacking ammonium sulfate and supplemented with the indicated amino acids. Growth was monitored in a 96-well plate using a SpectraMax i5 instrument for 24 h at 30°C.

### Generation of deletion mutants

A homologous recombination–based strategy was used to generate *C. glabrata* deletion mutants in both the BG2 and Bg14 backgrounds. *PUT1* and *PUT2* were disrupted using a hygromycin resistance cassette (hph), similar to the strategy used for *GCN2* and *GCN4* in a previous study (17). *PUT1* and *PUT2* deletion mutants were generated using the hph cassette derived from plasmid pAP599. For constructing deletion cassettes, the approximately 500 bp 5′ and 3′ untranslated regions (UTRs) of *CgPUT1* and *CgPUT2* were amplified from WT genomic DNA. These fragments were cloned on either side of the hph cassette in pAP599 using KpnI–HindIII (5′ UTR) and SacI–SpeI (3′ UTR) restriction sites. The *PUT3* deletion mutant was generated using an overlapping PCR strategy with a nourseothricin (NAT) resistance marker. Three PCR reactions were performed to obtain: (1) a 5′ UTR fragment fused to a short region homologous to the NAT marker, (2) a NAT fragment containing short homology to the PUT3 5′ UTR, and (3) a fused 5′ UTR–NAT product generated using fragments (1) and (2) as templates.

A similar approach was used to generate the 3′ UTR–NAT construct. The final overlapping PCR products containing the 5′ UTR–NAT and NAT–3′ UTR fragments were PCR-purified and transformed into *C. glabrata*. The linearized hygromycin and PUT3–NAT cassettes were transformed using the ethylene glycol–PEG-mediated transformation method, and transformants were selected on plates containing hygromycin (500 µg/ml) or nourseothricin (150 µg/ml). Successful gene disruptions were verified by diagnostic PCR using cassette- and backbone-specific primers. Complementation utilized uracil auxotrophy as a selectable marker in vectors pGRB2.2 or pCU-PDC1. The coding regions of *CgPUT1*, *CgPUT2*, *CgPUT3*, and *CgPUT4* were amplified from WT genomic DNA using Phusion polymerase. Amplicons were cloned downstream of either the PGK1 or PDC1 promoters in the respective plasmids. Luciferase reporter constructs for *PUT1*, *PUT2*, *PUT3* (full and half constructs), and *PUT4* were generated analogous to the *GCN4* luciferase reporter strategy (17). The 5′ UTR regions were amplified from genomic DNA using specific primer sequences and cloned into the SacI– BamHI sites of pGRB2.2, upstream of the firefly luciferase gene. The *PUT3* promoter lacking the GCN4-binding site was generated by overlapping PCR. Two template fragments devoid of the target nucleotide sequences were first amplified, followed by a final PCR to generate the full promoter with the deletion. All constructs were validated by restriction digestion and amplicon sequencing. The plasmids used in this study are listed in Supplementary Table S2, and the primers and restriction sites are listed in Supplementary Table S3.

### Polysome profiling

The WT *C. glabrata* strain was grown in SD medium to an OD_600_ of approximately 0.7–1 in a 500-ml culture. Untreated samples were harvested after incubation with 50 µg/ml cycloheximide (Sigma-Aldrich 01810) for 10 min. The remaining cultures were pelleted, washed, and transferred to SD medium lacking ammonium sulfate for proline utilization at 30 min, 120 min, and 240 min. Cycloheximide was added for the final 10 min before harvesting. Cells were centrifuged at 4°C and 4,000 rpm for 10 min. The pellet was washed once with the polysome lysis buffer and then resuspended in 500 µl of the same buffer. It was subsequently lysed mechanically using glass beads. Five OD_260_ units of whole cell extracts (WCEs) were layered on a sucrose gradient, and ultracentrifugation was performed. Subsequent steps, buffer compositions and instruments used were as described in a previous study (17). The polysome profile was analyzed, and the polysome/monosome ratio was determined from the AUC using ImageJ.

### Protein extraction and eIF2α phosphorylation assay

The overnight primary culture was used to inoculate the secondary culture in the minimal SD media. At the log phase, cultures with an OD_600_ of approximately 3 were harvested, serving as the untreated/control samples. The remaining cultures were washed and resuspended in SD medium lacking ammonium sulfate for proline utilization at the same time points (30 min, 120 min, and 240 min), after which cells with an OD_600_ of approximately 3 were collected. The WCEs were then prepared utilizing the TCA precipitation method (10% and 20% TCA), followed by resuspending the pellet in Tris and 2X Laemmli buffer and subsequent heat denaturation (17, 51). The prepared WCEs were resolved using SDS-PAGE, transferred to a nitrocellulose membrane, and probed with 1:4000 of Phospho-eIF2α (44-728G) antibody. Ponceau staining and GAPDH (ab22555) were used as the loading control.

### RNA extraction, cDNA synthesis, and quantitative real-time PCR

The RNA from WT and mutant strains was extracted from cells grown in SD medium at log phase, as well as in SD medium without ammonium sulfate and supplemented with amino acids. For each extraction, approximately 50 ml of the culture (OD_600_≈ 0.7–1) was harvested. The cells were washed with sterile DEPC-treated water, flash-frozen, and stored at -80°C. RNA was extracted using the hybrid protocol, which combines the classic acid–phenol extraction method with the Qiagen RNeasy kit (Cat. No. 74104), as described previously (17, 52). RNA was eluted in RNase-free water, and its concentration and purity were assessed by agarose gel electrophoresis and nanodrop analysis. One microgram of total RNA from each sample was used for cDNA synthesis, following the manufacturer’s instructions (Bio-Rad, USA: Cat. No. 1708891). Quantitative real-time PCR was performed using SYBR green master mix (Bio-Rad Cat. No. 1708891) in a reaction volume of 20 µl. The housekeeping genes encoding 5S rRNA, UBC13, and GAPDH were used for normalization.

### Luciferase assay

*Candida glabrata* Bg14 WT, *gcn2Δ*, *gcn4Δ*, and *put3Δ* cells were transformed with the luciferase reporter constructs harboring various proline pathway gene promoters. The strains were cultured in SD medium, and secondary cultures were inoculated and grown until the log phase (OD_600_ ≈ 0.7–1). Cells were then washed with sterile phosphate-buffered saline (PBS) and transferred to minimal SD media lacking ammonium sulfate. This was followed by induction with different amino acids for 30 min, 120 min, and 240 min, with the SD media control maintained throughout. After induction, the samples were adjusted to an OD_600_ of 1 in PBS, and the cells were flash-frozen and stored at -80°C until further analysis. Subsequently, luciferase activity was measured with the Promega kit (Cat. No. E6110) using a luminometer, and the Bradford assay was utilized to measure total protein content in the samples for normalization. The final values were plotted as relative luciferase units normalized per microgram of protein.

### Proline estimation

Proline levels were estimated using the CheKine™ Micro Proline (PRO) Assay Kit (Cat. No.: KTB1430), according to the manufacturer’s instructions with minor modifications. Log-phase cells (OD_600_ ≈ 0.7–1) grown in ammonium sulfate media were pelleted, washed, and subjected to proline utilization for 2 h. The cells were resuspended in the extraction buffer, bead-beaten, and processed further by boiling and centrifugation. The resulting supernatant was mixed with the assay buffer and chromogen processed with toluene extraction according to the kit’s protocol. The appropriately diluted supernatant was measured at 520 nm, alongside blank and positive controls. The positive control and the standard were prepared from the supplied powder. A standard curve was then generated (absorbance versus known proline concentration). Proline concentration was measured in µg/ml from the standard graph and normalized to the total protein content, as determined by the Bradford assay.

### Promoter analysis of *PUT3* for Gcn4 motif in pan- and non-*Candida* species

We identified *PUT3* orthologs across *Candida* spp. (*C. albicans*, *C. dubliniensis*, *C. glabrata*, *C. parapsilosis*, and *C. auris*) and non-*Candida* fungi (*S. cerevisiae*, *S. pombe*, *Aspergilus. fumigatus*, *C. neoformans*, and *Kluyveromyces. lactis*) using the *Candida* genome (CGD), National Center For Biotechnology Information (NCBI), and FungiDB databases. For each gene, the 1-kb upstream sequence of the annotated translation start codon (in correct gene orientation) was extracted and saved in FASTA format. These promoter sequences were scanned for occurrences of the *C. glabrata PUT3* motif TGAGTCAG, allowing up to two mismatches. The resulting representative 8-bp motifs were used to generate a cross-species motif logo using the ggseqlogo R package.

### Gcn4 and *PUT3* structure preparation

Gcn4 structure from *S. cerevisiae* was retrieved from the RCSB PDB with ID:1YSA (53). The *PUT3* DNA duplex was built as a 62-bp double-stranded DNA using 3DNA 2.0, corresponding to a 31-bp sequence (5’-TCGTTTCTC**TGAGTCAG**ACCTCTTTTGCTTC-3’) per strand (54).

Next, we performed energy minimization of both Gcn4 and *PUT3* using AmberTools22 (55). Both structures were energy-minimized in implicit solvent prior to docking. The protein was parameterized using the ff14SB force field, and the DNA was parameterized using the bsc1 force field, both in LEaP. Each structure underwent 4,000 steps of minimization (500 steps of steepest descent followed by 3,500 steps of conjugate gradient) in a Generalized Born implicit solvent model (igb = 5; 0.1 M salt; and 16 Å cutoff) to relieve initial steric clashes.

### Molecular docking

To determine the optimal conformation and orientation of *PUT3* when interacting with Gcn4, molecular docking was performed using the HDOCK server (56). Energy-minimized structures were used as inputs. HDOCK generated the top 10 predicted Gcn4–*PUT3* complexes. , The second model contained the largest number of interface residues with the TGAGTC motif of *PUT3* hence it was selected.

### ROS measurement

ROS levels in the *C. glabrata* (Bg14) WT/V, *put3Δ/*V, and *put3Δ/PUT3* strains were measured using H2DCFDA dye (ThermoFisher Scientific cat no-D399). This dye diffuses across the cell wall and is oxidized by intracellular ROS into a fluorescent molecule, DCF (Dichlorofluorescein), whose intensity correlates with ROS levels. Cells were grown in SD (ammonium sulfate) media with glucose or glycerol to log phase (OD_600_ ≈ 0.7–1), followed by 4 h of proline utilization in minimal media lacking ammonium sulfate with the respective carbon source. The cells from the ammonium sulfate and proline treatments on both carbon sources were washed, resuspended in PBS, and adjusted to an OD_600_ of approximately 1. Samples were incubated with 10 µM of H2DCFDA dye for 45 min at 30°C and 200 rpm in the dark. Controls included cells without dye, dye alone, and dye with H_2_O_2_, serving as the positive control. After treatment, cells were washed, resuspended in PBS, and split into two portions. One portion was used for fluorescence measurement (excitation: 485 nm; emission: 520 nm) using the Spectramax i5 plate reader, and the other was flash-frozen for total protein quantification by the Bradford assay. ROS levels, represented as relative fluorescence units (RFUs), were normalized to the total protein for each strain.

### Macrophage infection assay

Macrophage infection assays with WT and *put3Δ* strains were performed as described previously (16, 17, 57). This study utilized a monocyte cell line, THP-1 (a human acute monocytic leukemia cell line-derived macrophage). THP-1 cells were maintained in RPMI-1640 medium supplemented with antibiotics (Pen-Strep, Gibco) and heat-inactivated serum (FBS, Gibco, Cat No. A5256701) at 37°C with 5% CO₂. These cells were then converted into macrophages by treating them with 20 nM phorbol-12-myristate-13-acetate and seeding them at a density of 1×10^6^ per well in a 24-well plate. After 24 h, the medium was replaced with one without antibiotics to prepare the cells for infection. For the infection, overnight yeast cells were inoculated into a fresh medium, and at the log phase, the cells were washed and resuspended in PBS. The macrophages were then infected at an MOI of 1:10 (macrophage:yeast) for 2 h in triplicate. After the infection, all time point wells were cleared to remove extracellular yeast with PBS two or three times. Macrophages from the 2-h time point were then lysed with PBS–0.1% Triton X, serially diluted, and plated on YPD agar to assess phagocytosis. Afterwards, CFUs were recorded at the following time points—12 h, 36 h, and 42 h—using the same method to study intracellular survival. The colonies were counted after 2–3 d, and a graph of CFU/mL was plotted in the Prism software.

### Mice infection studies (disseminated candidiasis model)

All animal experiments were conducted at the Regional Centre for Biotechnology, Faridabad, India, in accordance with the guidelines of the CPCSEA. Procedures were designed to minimize animal suffering and were approved by the Institutional Animal Ethics Committee [RCB/IAEC/2022/131]. The female BalB/c mice (6–8 weeks old) were housed in well-ventilated cages under standard conditions in the animal facility. For the infection experiments, a secondary culture of yeast cells was prepared from an overnight culture. The cells were harvested at the log phase, washed with PBS, and adjusted to a concentration of 1×10⁸ cells. Mice (3–4 per strain) were injected intravenously with 1×10^8^ cells in 200 µl PBS . The infected animals were monitored every alternate day for clinical parameters, including body weight and temperature. After 7 d, mice were euthanized using CO_2_, and organs such as kidneys, liver, and spleen were aseptically harvested cleaned, weighed, and homogenized in PBS (58, 59). The organ homogenates were serially diluted and plated on YPD agar supplemented with penicillin and streptomycin, and plated in triplicate. The plates were then incubated at 30°C for 3–4 days. CFUs normalized to organ weight and expressed as CFU/g were plotted. Data from two independent experiments and a total of 6–8 mice were pooled, and statistical significance was assessed using a two-tailed *t*-test.

## ACKNOWLEDGEMENTS

The authors thank all members of Anil Thakur’s lab (RCB) for their valuable suggestions and assistance, especially in polysome profiling and mouse experiments. We are thankful to Dr. Rupinder Kaur for providing us with *C. glabrata* strains and plasmids. We are also grateful to Aman Srivastav for the thoughtful reading of the manuscript and his constructive suggestions. AR acknowledges ICMR for the fellowship.

## DATA AVAILABILITY STATEMENT

Supplementary Tables S1, S2, and S3 list the strains, plasmids, and primers used in this study, respectively. All relevant data are included in the manuscript and its supplementary materials. Additional data supporting the findings of this study are available from the corresponding author upon request.

## AUTHOR CONTRIBUTION

Conceptualization: AR, AT

Methodology: AR, SKG, AT

Investigation: AR, SKG, AT

Visualization: AR, SKG, AT

Supervision: AT

Writing—original draft: AR, AT

Writing—review & editing: AR, AT

## FUNDING

This work was supported by Anusandhan National Research Foundation (ANRF)/ Science and Engineering Research Board (SERB): EEQ/2022/000606, and RCB core.

## CONFLICTS OF INTEREST

The authors declare that there are no conflicts of interest

### Abbreviations

eIF2: Eukaryotic initiation factor 2
tRNA_i_^Met^: initiator tRNA
SD: Simple Dextrose Media
OD: Optical Density
AUC: Area under curve

## Notes

### Competing Interest Statement

The authors have declared no competing interest.

